# Sensory processing dysregulations as reliable translational biomarkers in *SYNGAP1* haploinsufficiency

**DOI:** 10.1101/2021.04.28.441866

**Authors:** Maria Isabel Carreño-Muñoz, Bidisha Chattopadhyaya, Kristian Agbogba, Valérie Côté, Siyan Wang, Maxime Lévesque, Massimo Avoli, Jacques L. Michaud, Sarah Lippé, Graziella Di Cristo

## Abstract

Amongst the numerous genes associated with intellectual disability, *SYNGAP1* stands out for its frequency and penetrance of loss-of-function variants found in patients, as well as the wide range of co-morbid disorders associated with its mutation. Most studies exploring the pathophysiological alterations caused by *Syngap1* haploinsufficiency in mouse models have focused on cognitive problems and epilepsy, however whether and to what extent sensory perception and processing are altered by *Syngap1* haploinsufficiency is less clear. By performing EEG recordings in awake mice, we identified specific alterations in multiple aspects of auditory and visual processing, including increased baseline gamma oscillation power, increased theta/gamma phase amplitude coupling following stimulus presentation and abnormal neural entrainment in response to different sensory modality-specific frequencies. We also report lack of habituation to repetitive auditory stimuli and abnormal deviant sound detection. Interestingly, we found that most of these alterations are present in human patients as well, thus making them strong candidates as translational biomarkers of sensory-processing alterations associated with *SYNGAP1/Syngap1* haploinsufficiency.

## Introduction

*SYNGAP1* haploinsufficiency-related intellectual disability (*SYNGAP1*-ID) is characterized by moderate to severe developmental delay or intellectual disability (ID), generalized epilepsy (>80%), autism spectrum disorder (ASD) and other behavioral abnormalities including severely delayed or absent language development, stereotypic behaviors, poor social development, impulsivity, inattention, aggressive behavior, elevated pain threshold and sleep disorders (>50%)^1–8^. To date more than 700 individuals with *SYNGAP1*-ID have been reported worldwide through genetic sequencing and this number is rising rapidly due to more widespread genetic testing. The majority of affected individuals carry *de novo* nonsense or frameshift variants that likely result in haploinsufficiency. *SYNGAP1* haploinsufficiency may explain up to 1% of non-syndromic ID cases^2, 3, 9, 10^, currently making it one of the most frequent causes of genetically-defined childhood brain disorders.

Using rodent models both *in vitro* and *in vivo*, research over the last twenty years has uncovered the critical roles played by SYNGAP1, the Ras GTPase-activating protein coded by the *Syngap1* gene, in neuronal physiology, brain development, and cognition (recently reviewed by Gamache and collaborators^11^). Critical to our understanding of SYNGAP1 function in the context of neurodevelopmental disorders, it has been proposed that *Syngap1* heterozygous knockout mice, which have ~50% reduction in SYNGAP1 protein levels^12^, offer both construct and face validity for *SYNGAP1*-ID^13^. Several studies have attempted, and succeeded, to rescue specific behavioral and neurological phenotypes in these mice^14, 15^. These exciting results deepen our neurobiological understanding of SYNGAP1 function in normal and haploinsufficient conditions. Nevertheless, a critical, and still missing, step for the successful development of therapies for *SYNGAP1*-ID is the identification of robust, replicable and translatable biomarkers in humans and animal models.

Electroencephalography (EEG) recordings and sensory-evoked event-related potentials are particularly well suited for the investigation of neurodevelopmental disorders^16^. Since EEG has a high temporal resolution, sensory-evoked related potentials can provide detailed brain-related readouts of cognitive processes. EEG is also non-invasive, portable, inexpensive, and can assess brain activity through passive tasks, and it thereby not dependent on patient’s active participation.

Sensory-evoked potentials exhibit distinctive temporal patterns of activity in specific brain regions that are replicable across trials and between typical individuals^16^. The reliability of EEG as an outcome measure in clinical trials is currently under investigation. Here, we assess whether EEG can be used as a translational (human-mice) investigation tool in *SYNGAP1-ID*.

Sensory-evoked potentials induced by a discrete sensory stimulus, usually visual or auditory, are currently being investigated as biomarkers of sensory processing impairment in neurodevelopmental disorders^16–20^. In fact, recent studies suggest that sensory-processing impairments are very frequent in neurodevelopmental disorders; they significantly affect social adaptation and behavioural preferences, and limit quality of life^19, 21^. A recent study reported abnormal tactile processing in individuals with *SYNGAP1*-ID, and severe impairments in somatosensory cortex physiology in *Syngap1* haploinsufficient mice^22^. Several parallel studies on the neurobiological alterations caused by *SYNGAP1/Syngap1* haploinsufficiency has focused on epileptic activity and cognitive problems. However, whether sensory processing, in particular auditory and visual processing, is impaired in both *SYNGAP1*-ID individuals and *Syngap1* haploinsufficient mice is as yet unexlored.

Here, we performed and analyzed EEG recording following different visual and auditory stimulation protocols in awake *Syngap1^+/-^* (haploinsufficient) mice and control littermates of both sexes. We further performed the same analysis in a cohort of patients with *SYNGAP1*-ID and of neurotypic age-matched individuals, who underwent EEG recording following both visual and auditory stimulations. We identified significantly increased gamma oscillation power at baseline consistent across different cortical regions, lack of habituation to repetitive auditory stimuli and increased coupling of theta and gamma oscillations following both visual and auditory stimulations as robust and translational sensory-processing alterations associated with *SYNGAP1/Syngap1* haploinsufficiency.

## Material and Methods

### EEG recordings in mice

#### Animals

Experimental procedures involving mice were approved by the Comité Institutionnel de Bonnes Pratiques Animales en Recherche (CIBPAR) of the Research Center of Sainte-Justine Hospital in accordance with the principles published by the Canadian Council on Animal Care. *Syngap1^+/−^* mice were kindly provided by Dr Seth Grant (Edinburgh University, United Kingdom)^23^. These carry a deletion of exons 8-10 of the *Syngap1* gene, which encode for the C2 and GAP domains^23^. They were maintained as a heterozygous line onto a C57Bl/6 background. Twenty-two adult *Syngap1^+/−^* mice (12 females and 10 males) were used in this study. Mice were implanted at postnatal day (P)70-90 and recorded between P80 and P120. All animals were housed with 2-5 mice/cage from weaning until surgery, after which they were housed singularly for the duration of the experiment. Animals were kept on a 12h/12h light/dark cycle, provided with nesting material and food and water *ad libitum*. All experiments were performed during the light period under constant mild luminosity (60 Lux). We recorded mice of both sexes. We checked and noted the exact stage of females’ estrous cycle after each recording session, by histological inspection of vaginal smear^24^. Here, we analyzed EEG only for females in the pro-estrus/estrus phase, with the exception of inter-ictal spike analysis for which we pulled together data from mice in either diestrus/metestrus or pro-estrus/estrus recordings.

#### Surgery

Cranial surgery was performed under deep anesthesia, induced by an intraperitoneal injection of ketamine/xylazine cocktail. A craniotomy was performed at the coordinates described below on the right side of the brains and custom-made clusters of four to six 50μm-diameter-insulated-tungsten wires were implanted. In each cluster, wires tips were 50 to 150 μm apart. A custom-designed and 3D printed microdrive-like scaffold held the 3 electrode clusters at fixed positions (Supplementary Figure 1), thus allowing the simultaneous implants of all 3 and reducing surgery time.

Data was acquired with a custom-made headstage using Intan Technologies’ RHD Electrophysiology Amplifier chip. A ground-and-reference wire was also gently introduced in the contralateral frontal lobe and a 2cm carbon fiber bar was placed on the back of the skull. The microdrive apparatus, the carbon fiber bar and the ground-and-reference wire were then daubed with dental acrylic to encase the electrode-microdrive assembly and anchor it to the skull. For one week after surgery, animals were treated with Metacam and the skin around the implant was disinfected daily.

Electrode clusters were implanted at the following coordinates: Primary Auditory Cortex (A1), - 2.5 mm posterior to Bregma, 3.9 mm lateral to the midline and 0.8 mm deep; Primary Visual Cortex (V1), −4 mm posterior to Bregma and 2 mm lateral to the midline; Posterior Parietal Cortex (PPC), −2 mm posterior to Bregma and 1.7 mm lateral to the midline. Clusters placed in V1 and PPC targeted both supra and infragranular layers, those placed in A1 targeted layer 5. PPC cluster also included 2 longer (1.2 and 1.3 mm) wires reaching the dorsal region of the hippocampus (CA1 pyramidal layer and radiatum). Correct depth in CA1 was confirmed by the presence of ripples, which appeared on the recordings when the electrode reached the pyramidal layer. For this study, we analyzed only signals from cortical layer 5 and hippocampal radiatum layer.

#### EEG recording

EEG recordings were performed using an open-ephys GUI platform (https://open-ephys.org/). Custom designed auditory and visual trigger generator devices sent triggers directly to the recording system assuring the time precision of the stimuli presentation. Starting one week after surgery, animals were habituated daily for 4 days to an open field (15-20’ each time), before being recorded during auditory stimulation protocols from day 5. We used a dimly illuminated open field environment (45×35cm), surrounded by 60cm-high transparent walls, and equipped with video monitoring. The walls were sprayed and wiped clean with 70% ethanol 30 minutes before the introduction of each animal. Two loudspeakers were placed ~10 cm from the walls of the open field. Baseline activity was recorded at the beginning of each session for a period of 10 minutes while mice were freely exploring the open field. EEG recoding during auditory stimulations were conducted for 20-30 minutes, a total of 4 times, on alternate days. Mice were then habituated for 7-10 days to the head-fixed system situated on air-supported ball treadmill. This custom-made apparatus allowed the animal to run using a larger range of directions than a classical treadmill. Habituation period was progressively longer, starting from 5 up to 30’ daily. Once mice were habituated to stay on the ball for 30 minutes, they were recorded during visual stimulation protocols. Each recording session lasted from 15-30 minutes.

#### Auditory stimulations protocols

All stimulations were presented at an intensity of 70 dB. For pure tone stimulation, we used 100 ms long 5KHz and 10KHz pure tones presented with a random inter-stimuli interval of 2 to 3s. Each pure tone was presented 60 times using a random order. The full protocol lasted 5 minutes. For the habituation/oddball paradigm, we used sequences of 5-9 repetitive (standard, at 5KHz) sounds followed by a deviant sound (at 10KHz). Sequences were presented in a pre-established pseudorandom order. Sounds lasted 70ms and were presented with a 1s inter-stimulus-interval. The full protocol lasted 10 minutes. For Auditory Steady State stimulation, we used 10 and 40 Hz click trains (1s duration and 2.5 inter trial interval) presented with alternating frequencies. Each click was a sound at 5KHz lasting 5ms. This protocol lasted for 5 minutes. Finally, we used a Chirp stimulation, adapted from^25^. Briefly, we used a 2s 5KHz tone whose amplitude was modulated by a sinusoid whose frequency increases (up chirp) or decrease (down chirp) linearly in the 1-100Hz. To avoid onset contaminating phase locking to the amplitude modulation of the chirp, the stimulus was ramped in sound level from 0-100% of its original intensity over 1s, which then transitioned into chirp modulation sequence mentioned previously. Inter-trial intervals were random in a range from 2-4s. Up and down chirp were presented 50 times each, in alternation. Both directions of modulation were tested to ensure any frequency specific effects were not due to the frequency transition history within the stimulus. The interval between each train was randomly varied between 1 and 1.5s. The total duration of this protocol was 15 minutes.

#### Visual stimulation protocols

All visual stimulations were presented using a 27-inch monitor with a resolution of 2560 x 1440 pixels and a refresh rate of 60Hz. The screen was placed 50 cm from the animal nose. We used alternating grating, horizontal black and white sinusoidal bars alternating at 1Hz, with fixed spatial frequency of 0.05 cycles/degree and 100% contrast^17^. Stimuli duration was 2s, presented at 2.5s inter-stimulus-interval. The whole protocol lasted 6 minutes. In addition, for Visual Steady State stimulation, we used flickering light flashes at different frequency (8, 15 and 30Hz) presented in trains of 1.2s, with 2s inter-stimulus-interval. This protocol lasted 15 minutes.

### EEG recordings in humans

#### Participants

We analyzed the EEG recordings from *SYNGAP1*-ID and neurotypical participants reported in^26^. Experimental procedures involving humans’ subjects were approved by the ethics, scientific and administrative committees of CHU Sainte-Justine. Written informed consent from parents or caregivers and participants’ assent were obtained before study participation. Data were collected from 8 participants diagnosed clinically with *SYNGAP1* haploinsufficiency and 49 age-matched neurotypical (NT) control subjects. Three to seventeen years old participants of both sexes were recruited. Only NT subjects without a family history of neurodevelopmental disorders such as ID or ASD were recruited. Exclusion criteria for all groups consisted of having undergone neurosurgery and/or having vision or hearing problems. Comprehensive clinical phenotype, inclusion criteria and pharmacological treatment of the clinical group is reported in^26^.

#### EEG recordings and sensory stimulation protocols

Recordings are described in^26^ and briefly summarized below. EEG recordings took place in an electrically shielded and sound-attenuated room in CHU Sainte-Justine. The recording was carried out using Geodesics 128 electrode nets (Electrical Geodesics System Inc., Eugene, OR, USA). Data was acquired using an EGI Net Amp 300 amplifier and stored on a G4 Macintosh computer with NetStation EEG Software (Version 4.5.4). The EEG procedure was heavily adapted to the clinical populations in order to increase acceptance. Pictograms and videos were used to prepare participants before coming in for the evaluation and storytelling was used to make the wearing of the EEG net more desirable. A movie was shown during the net installation to increase participants’ collaboration. Lights were turned off during the recording session. Auditory stimuli were presented via two lateral speakers (BX5a, M-Audio^®^ BX studio, Cumberland, RI, USA) at a distance of 30 centimeters from participants’ ears and visual stimuli were presented on a computer monitor, at a viewing distance of 60 cm. Stimuli were generated by a Dell Optiplex 790 PC using E-Prime 2.0 (Psychology Software Tools, Inc., Pittsburgh, PA, USA). EEG was recorded at a sample rate of 1000Hz using the vertex electrode as online reference. Impedances were kept below 40 kΩ^27^. During the white noise task and the resting state recording, a silenced movie without subtitles was presented to ensure maximal collaboration. Participants were told to watch the movie and listen passively to the sounds.

We selected three regions of interest (ROI) according to literature of sensory processing in EEG. Auditory processing is reported to peak in vertex and midline electrodes^28, 29^, thus we selected Midline central (Cz) surrounding electrodes as ROI for our analyses on auditory perception, as the equivalent to the Primary Auditory Cortex in mice (supplementary figure 1). Cz ROI is composed of the five electrodes surrounding Cz. Visual perception is recorded at the occipital region. Occipital ROI is composed of Oz electrode in addition to four surrounding electrodes. A third ROI corresponding to the Posterior Parietal Cortex was used by grouping Pz electrode and the five surrounding electrodes as indicated in supplementary figure 1. Each ROI was defined as the average of the corresponding cluster of electrodes mentioned above. Finally, Cz electrode was used as the digital reference and COM electrode as the analog reference.

Baseline (resting) activity was analyzed during the first three minutes of the recording session. During this time, participants were watching a muted film to engage their participation and encourage immobility. The following sensory stimulation protocols were used. White Noise: Auditory stimulation protocol consisted of a 24 dB/ octave white noise burst presented 150 times. Each stimulus lasted 50ms with an inter-stimulus-interval varying between 1200 and 1400ms to avoid habituation. Total task duration was about 4 minutes. Checkerboard: Visual stimulation was a pattern-reversal black and white checkerboard with a spatial resolution set to 2*2cm and a luminance of 40cd/m2. Stimulation protocol lasted for 4 minutes with a total of 200 repetitions.

Pattern reversals were done every 500ms for the whole duration of the protocol with no inter stimulus interval. A fixation cross was only presented at the beginning of the task in order to attract the subject’s visual attention. Habituation Paradigm: audio-visual speech stimuli were used in order to maximize participants’ attention^26, 30^. The task had in total 96 sequences of four phonemes /aaai/ and /aaaa/ (75% /aaai/ and 25% /aaaa/). Male or female faces are presented with their matched phoneme track four times (aaai or aaaa) then a new sequence start with the other sex. Here analysis was conducted only on the auditory responses elicited by the first and second /a/ to assess the most prominent habituation response. The first presentation of /a/ is referred here as S1 and the second one as S2.

### EEG analysis in mice and humans

#### EEG pre-processing

Mice EEG signal was downsampled to 2000 Hz. Wideband filter (0.5-1000Hz) was applied for ictal spike analysis and a narrower passband filter (0.5-150Hz) was applied for the rest of analysis. A notch filter (59.5–60.5 Hz) was also applied to remove residual 60 Hz power-line noise contamination. Data were then segmented into periods of different length depending on the stimulation protocol: 1400 ms (500 ms pre- and 900 ms post-stimulus onset) for auditory oddball protocol, 2500 ms (1000 ms pre and 1500 ms post-stimulus onset) for pure tone stimulation, 3000 ms (1000 ms pre and 2000 ms post-stimulus onset) for alternating grating and 4000 ms (1000 ms pre and 3000 ms post-stimulus onset) for both visual and auditory steady state. Trials containing per sample segments with an intra-channel average higher than 4 times the total trial standard deviation, were tagged, visually inspected, and removed. Only sessions containing more than 50 clean trials were kept for further analyses.

Humans EEG signal pre-processing and segmentation were carried out using BrainVision Analyser Software (Brain Products, Munich, Germany). EEG raw data was filtered with a lower bound of 0.5 Hz and a high cut-off at 100Hz during data acquisition. A selective 60Hz notch filter was applied. Twenty-eight electrodes around the neck and the face were excluded from analysis since they contained muscular artifacts and thus were not useful for further analyses. Blink artefacts and remaining muscular artefacts were removed using a semi-automatic independent component analysis (ICA)^31^. Data were re-referenced to an average reference. For ERP analysis, data was segmented from −200 to 500 ms relative to stimulus onset and the signal was baseline corrected from −150 to 0 ms. For inter-trial coherence (ITC) and time-frequency spectrograms (TFS), epochs were created −800 to 1000 ms relative to stimulus onset. TFS were baseline corrected from −500 to 0 ms.

The semi-automatic artefact rejection procedure marked segments with voltage exceeding two standard deviations or −200 to 200 uV of amplitude range, which were rejected during the following manual artefact rejection in addition to other segments that were judged noisy. Following artifact rejection, for the white noise paradigm, an average of 139.91/ 150 (SD = 11.85) epochs were kept for final analysis in control participants, and 105/ 150 (SD = 35.94) for *SYNGAP1-ID* participants. For the habituation paradigm, an average of 64.22 / 80 (SD = 14.52) epochs were kept for control and an average of 52.17/ 80 (SD = 21.19) for *SYNGAP1*-ID participants for both S1 and S2. For the checkerboard paradigm, an average of 126.15/ 200 (SD = 43.48) epochs were kept for the control participants and an average of 81.83/ 200 (SD = 35.88) epochs for *SYNGAP1-ID* participants. All epochs were ictal free.

#### Data Analysis

Mice and human data were analyzed using the same algorithms. None of the recorded animals were excluded from the final analysis. Signal analysis and quantification was performed using custom MATLAB (The Mathworks Inc., Natik, MA, USA) code, available upon request.

##### Interictal spikes detection in mice

For each animal, a 2 hours-long epoch was acquired between 10 AM and 4 PM. We then analyzed the first 1 hour of EEG recording performed when mice were freely moving and awake in the open box after the habituation period. We compared rates of interictal spikes and rates of interictal spikes with high frequency oscillations (HFOs) between groups. Processed data were then exported to Matlab R2020a (The Mathworks, MA, USA) and analyzed off-line. Interictal spikes were detected based on threshold crossings (mean and standard deviation), calculated over the entire period. Events above 4 SD were considered as potential interictal spikes. Every detected event was then analysed visually, and false positives caused by movement artefacts were removed manually. Only events detected during periods of immobility (<2.5cm/s) are shown in Figure 1 and Supplemental Figure 2.

**Figure 1.**
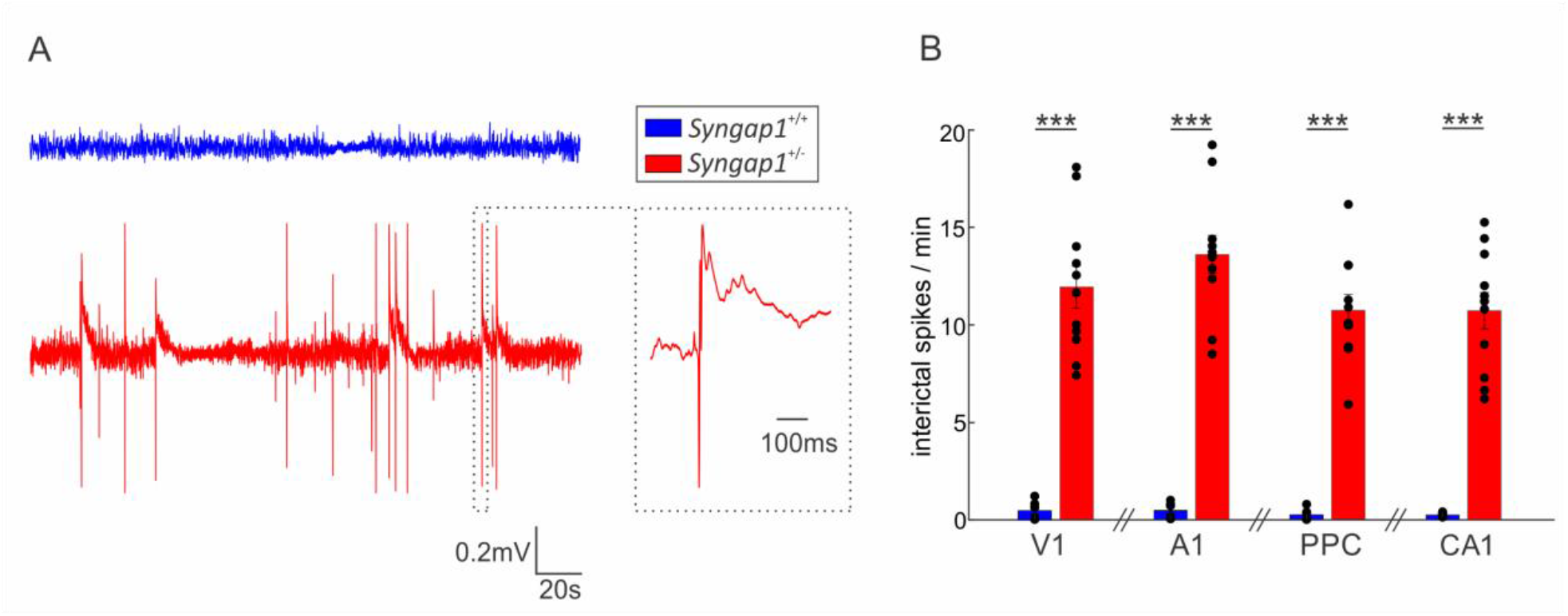
*Syngap1* haploinsufficient mice show increased inter-ictal spike activity. **A:** Example of EEG trace recorded from of a *Syngap1^+/+^* (blue) and *Syngap1^+/-^* (red) mouse during 2 minutes of immobility/sleep. On the right, interictal spike on an expanded time scale. **B:** Bar plots show the total number of interictal spikes detected in V1 (t-test t(15)=-7.6610, p<0.0001), A1 (t-test; t(15)=-9.9019, p<0.0001), PPC (t-test; t(15)=-9.3477, p<0.0001) and CA1 (t-test; t(15)=-7.4481, p<0.0001). *** indicates p<0.0001. Bar graphs represent mean±SEM. *Syngap1^+/+^* n=7; *Syngap1^+/-^* n=11 mice.

##### Visual Evoked Potential (VEP) and Auditory Evoked Potential (AEP) in mice and humans

EEG signal was low pass filtered to 150 Hz, baseline corrected to the mean voltage of the 150 ms prior stimulus onset and averaged over trial. ERP components baseline-to-peak N1 and P1 were analyzed from AEP and VEP. N1 amplitude was automatically detected by subtracting the minimum voltage (negative peak) within a 10- to 80-millisecond time window after stimulus onset to the averaged baseline value. P1 amplitude was extracted by subtracting the maximum voltage (positive peak) within an 80- to 150-millisecond time window after stimulus onset to the averaged baseline value. Analyses were done using Fieldtrip toolbox v 202009^32^.

##### Habituation and Mismatch negativity (MMN)

For quantifying habituation in mice and humans, we calculated the ratio between the amplitudes of the first standard sound (S1) and the second (S2). These ratios were then converted to logarithmic based 10 scale. Mismatch negativity was calculated by subtracting the EEG trace of the standard sound to the EEG trace of the deviant sound. The obtained pattern was then used to detect N1 and P1 amplitudes in each subject.

##### Power Spectral Density in mice and humans

Resting state data segments were selected with 50% overlap and a hamming window was applied in order to reduce the amplitudes of the discontinuities at the boundaries of the epochs. Fast Fourier transform was applied using the fft() function available in MATLAB.

##### Time Frequency Spectrogram (TFS) in mice and humans

TFS analysis allows to explore the power of different frequency bands over time^33,34^. We used complex Morlet’s wavelet transformation^35^ to provide power maps in time and frequency domains. The simplified mathematical expression for applying this specific wavelet convolution is:

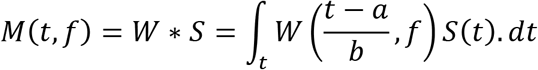

*M*(*t, f*) is a matrix of complex values (vectors) for a given time (t) and frequency (f). *S* is the signal as a function of time (t) and W corresponds to Morlet’s wavelet which is a complex exponential (Fourier) with a Gaussian envelope which undergoes a series of translations (a) and dilations (b) dependently of the frequency (f). Wavelet cycles increase linearly with frequency, beginning at 2Hz [2,0.5]. Baseline correction was applied using the 500 ms prior the stimuli onset. This analysis was performed using EEGLAB toolbox (v.13.6.5b) (La Jolla, CA, USA)^36^, in particular using the function *newtimef*.

##### Inter-trial coherence (ITC)

Analogous to PLV (Phase-locking value), ITC allows the assessment of the strength of phase coherence across trials in temporal and spectral domains^34^. The ITC computation uses only the phase of the complex values given by Morlet’s wavelet transform. ITC measures phase coupling across trials at all latencies and frequencies and is defined by:

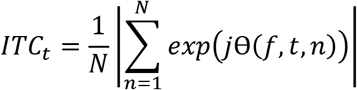

where *j*θ(*f,t,n*) represents the phase for a given frequency (f), time point (t) and trial (n). The obtained values are always defined between 0 and 1. Phase-locking values close to 1 indicate strong inter trial phase-locking, thus representing evoked activity while scores closer to 0 indicate a high inter trial phase variability^33, 37^.

##### Phase Amplitude Coupling (PAC)

PAC analysis allows the evaluation of nesting oscillations, where the phase of the lower frequency (here called fp) modulates the amplitude of the higher frequency (fa). The bandwidth of the filter used to obtain ‘fp’ and ‘fa’ is a crucial parameter in calculating PAC^38^. The filters for extracting ‘fa’ need to be wide enough to capture the center frequency ± the modulating fp. If this condition is not met, then PAC may not be detected even if present^39^. We therefore decided to use a wide enough bandwidth for both ‘fa’ and ‘fb‘. The coupling between ‘fp’ and ‘fa’ was quantified using the mean vector length modulation index (MVL-MI)^38^ as it has been suggested to be more sensitive to coupling strength and width compared to other methods^39^. This approach estimates PAC from a signal with length N, by combining phase (ϕ) and amplitude information to create a complex-valued signal.

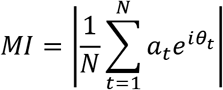

where N is the total number of data points, t is each time point, at is the amplitude of ‘fa’ at the time point t and θ_t_ is the phase angle of ‘fp’ at time point t.

To evaluate whether the observed modulation index actually differs from what would be expected by chance, surrogate analysis needs to be performed. To do so, surrogate data, are created by shuffling trial and phase-carrying information (500 surrogates), to normalize MI-values. These random permutations create a new time series with broken temporal relationships between the phase and amplitude information. Then MI is estimated again using the shuffled time series to obtain the null distribution of surrogate modulation index values. A normalized modulation index (MI_norm_) is then obtained as a *z*-score:

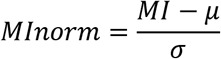

where μ and σ are the mean and standard deviation obtained from the null distribution. A Bonferroni corrected t-test was then performed and all MI-values exceeding a 5% significance threshold were grouped into clusters for further statistical analysis by group. This analysis was performed using PACmeg code provided by^40^.

### Statistical analysis

Data were systematically tested for Normal distribution, with the Lilliefors test, which is a modification of the Kolmogorov-Smirnov test recommended for small sample sizes^41^. Homoscedasticity was assessed using the Levene’s test. Differences between two groups of normally distributed data with homogeneous variances were analyzed using parametric student’s t-test, while not normally distributed data were analyzed with Wilcoxon rank sum test. To evaluate sex effect, 2-ways ANOVA (F) test were performed. When no sex effect was observed, main effect (genotype) was analyzed using Tukey *posthoc* test. For the auditory habituation test S2/S1 ratios were converted to logarithmic based 10 scale because distributions were not homogeneous. Paired sample test was performed using paired T-test when normality was fulfilled, and Wilcoxon signed rank test when it was not. Significance level was set to 5% (*p* = 0.05). Results were considered significant for values of p<0.05. Data are presented as mean ± standard error of mean.

### Data availability

All data are available upon reasonable request.

## Results

### *Syngap1^+/-^* mice show ictal activity in multiple cortical regions during immobility

Most *SYNGAP1*-ID patients have epilepsy^2–8, 42^. Interestingly, previous studies reported increased frequency of interictal spike events and occasional electrographic seizures in two different mouse models of *Syngap1* haploinsufficiency^14, 15, 43^, which appeared to be most frequent during non-rapid eye movement sleep^14, 15^. Interictal spikes are pathological electrical events that reflect seizure susceptibility in epileptic patients^44^. However, since EEG studies had not been previously performed in the *Syngap1* heterozygous mice line used in our study, if and whether they show interictal spike was unknown. Therefore, as first step we performed video-EEG recordings on freely moving mice in an open field and analyzed interictal spikes during immobility. We observed frequent interictal spikes, both sparse and in clusters, in all the recorded *Syngap1^+/-^* mice independent of the recorded regions (Fig.1). During the recordings, we observed only one electrographic seizure in one *Syngap1^+/-^* mouse, which was not accompanied by clear motor manifestations. We further compared the frequency of interictal spikes in males and females and found that it was independent of the sex (Supplemental figure 2). These results demonstrate that high-amplitude, pathological interictal spikes, in particular during immobility, are a reproducible biomarker across different *Syngap1* haploinsuffícient mouse lines^14, 15^.

### Increased baseline gamma power is a common phenotype in both *Syngap1^+/-^* mice and *SYNGAP1*-ID patients

Neural oscillations in the gamma band (35-100Hz) are associated with sensory processing and cognition^45^. Remarkably, abnormal gamma oscillations have been reported in a variety of neuropsychiatric and neurodevelopmental disorders, including ASD^46–49^ and schizophrenia^50–52^. Here, we performed power density spectra analysis on 3-minutes EEG periods during baseline activity (mice were freely exploring the Open Field but no stimulus was presented) in *Syngap1^+/+^* and *Syngap1^+/-^* mice and quantified mean values for beta (14-30 Hz), low gamma (35-55 Hz) and high gamma (65-100 Hz). *Syngap1^+/-^* mice showed a statistically significant increase in low gamma and high gamma power compared to wild-type littermates, which was consistent in all the recorded regions (Figure 2A, B). Both females and males *Syngap1^+/-^* mice showed increased gamma power, although appeared to be more pronounced in males (Supplementary figure 3).

**Figure 2.**
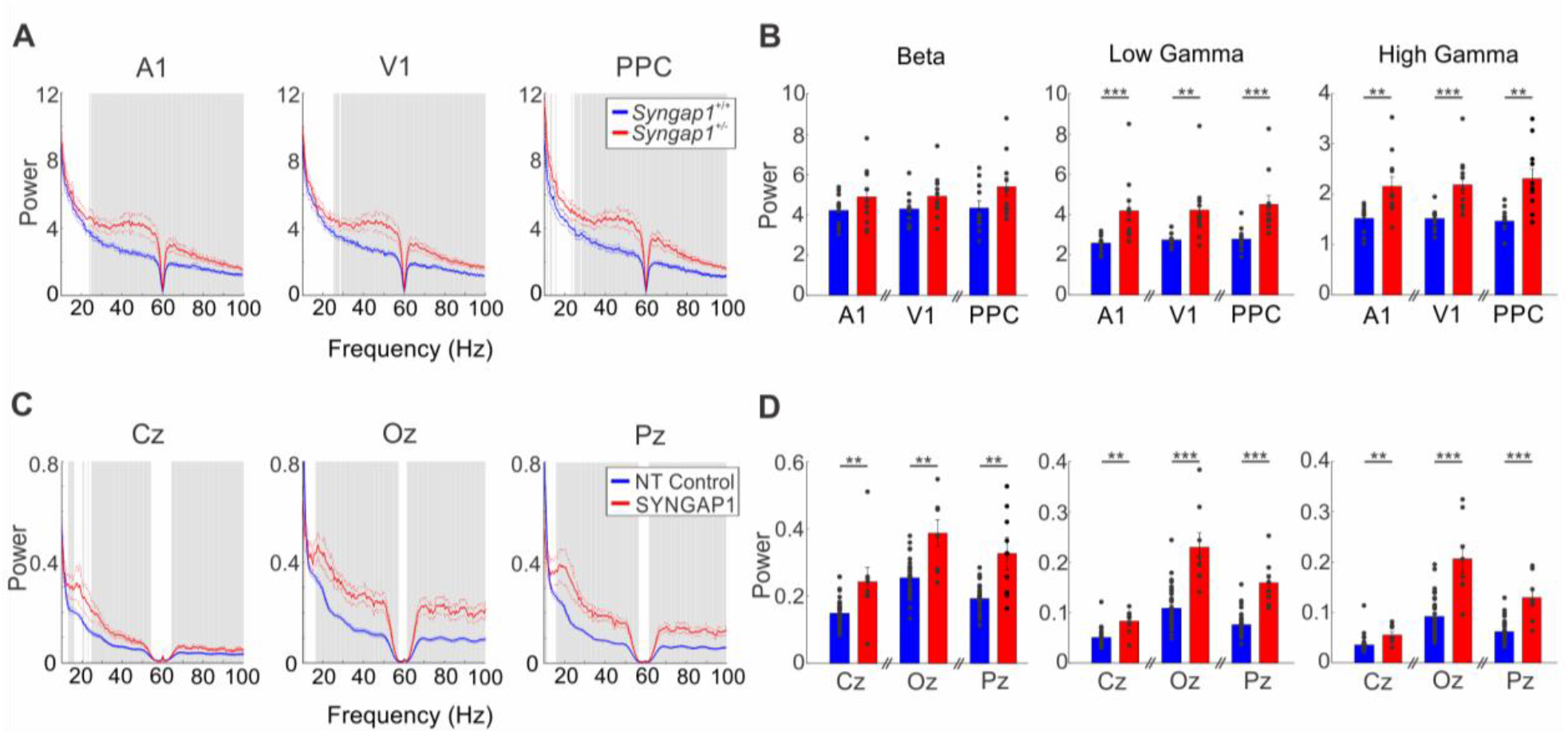
*Syngap1/SYNGAP1* haploinsufficiency correlates with increased baseline gamma power in both mice and humans during baseline state. **A, C**: Power Density Spectra of 3-minutes baseline state (non-stimulus) EEG signal from A1/Cz, V1/Oz and PPC/Pz, in mice (**A**) and humans (**C**). Gray bars represent significant differences (t-test, p<0.05). **B,D**, Bar plots shows mean powers±SEM of the **beta band** in mice (A1: Wilcoxon rank sum test; z=-1.0506, p=0.2934; V1 : Wilcoxon rank sum test; z=-1.5103, p=0.1310; PPC Wilcoxon rank sum test; z=-1.9043, p=0.0569) and humans (Cz: Wilcoxon rank sum test; z=-2.7387, p=0.0062; Oz : Wilcoxon rank sum test; z=-3.0741, p=0.0021; Pz Wilcoxon rank sum test; z=-2.7308, p=0.0063), **low gamma band** in mice (A1: Wilcoxon rank sum test; z=-3.3489, p=0.0008; V1 : Wilcoxon rank sum test; z=-3.2176, p=0.0013; PPC Wilcoxon rank sum test; z=-3.4802, p=0.0005) and humans (Cz: Wilcoxon rank sum test; z=-2.9949, p=0.0027; Oz : Wilcoxon rank sum test; z=-3.9480, p=0.00007; Pz Wilcoxon rank sum test; z=-4.0728, p=0.00004), and **high gamma band** in mice (A1: Wilcoxon rank sum test; z=-3.0206, p=0.0025; V1 : Wilcoxon rank sum test; z=-3.4802, p=0.0005; PPC Wilcoxon rank sum test; z=-3.2176, p=0.0013) and humans (Cz: Wilcoxon rank sum test; z=-3.1551, p=0.0016; Oz : Wilcoxon rank sum test; z=-3.7295, p=0.00019; Pz Wilcoxon rank sum test; z=-3.6983, p=0.00021). Note significantly increased power of low and high gamma band (** p<0.01; *** p<0.001) in mice and humans. Number of mice: *Syngap1^+/+^* n=11; *Syngap1^+/-^* n=11; number of human participants: NT Controls n=35; *SYNGAP1*-ID n=8.

Further, to investigate whether *SYNGAP1* haploinsufficiency is also associated with increased baseline gamma power in humans, we performed the same analysis as we did in mice using EEGs previously recorded from *SYNGAP1*-ID individuals^26^. Similar to what was observed in the mouse model, we found a statistically significant increase in both low and high gamma bands, consistent across all regions of interested (Figure 2C, D), during baseline activity. We further observed a significant increase in beta power in *SYNGAP1*-ID patients which was not present in *Syngap1^+/-^* mice (Figure 2B, D). These data suggest that increased power of the baseline (non-stimulus) gamma band is a robust and translational biomarker for *Syngap1/SYNGAP1* haploinsufficiency.

### *Syngap1/SYNGAP1* haploinsufficiency in associated with excessive theta/gamma phase-amplitude coupling following visual and auditory stimulations

Sensory-evoked potentials induced by simple sensory stimulus, in particular visual-evoked potentials (VEPs) and auditory-evoked potentials (AEPs) described by the waveform of EEG activity in the first 500 ms after stimulus presentation, are the cortical responses most commonly used to study sensory perception in both humans and mouse models^16^. They been proposed as biomarkers for several neurodevelopmental disorders, including ASD^16, 17, 20, 53, 54^. Here, we analysed VEP and AEP amplitudes in *Syngap1^+/-^* mice and control littermates (Figure 3). Similar to published reports from Rett’s syndrome patients and mouse models^17^, *Syngap1^+/-^* mice exhibited reduced amplitude of the P1 component of the VEPs compared to wild-type mice (Figure 3A). On the other hand, AEPs analysis showed increased N1 amplitude but no change in P1 in the *Syngap1^+/-^* mice (Figure 3C). For both VEPs and AEPs, we observed a significant effect of the *Syngap1* genotype, but not sex-related differences, and no statistical significant interaction was detected between these two factors (Supplementary Figure 4).

**Figure 3.**
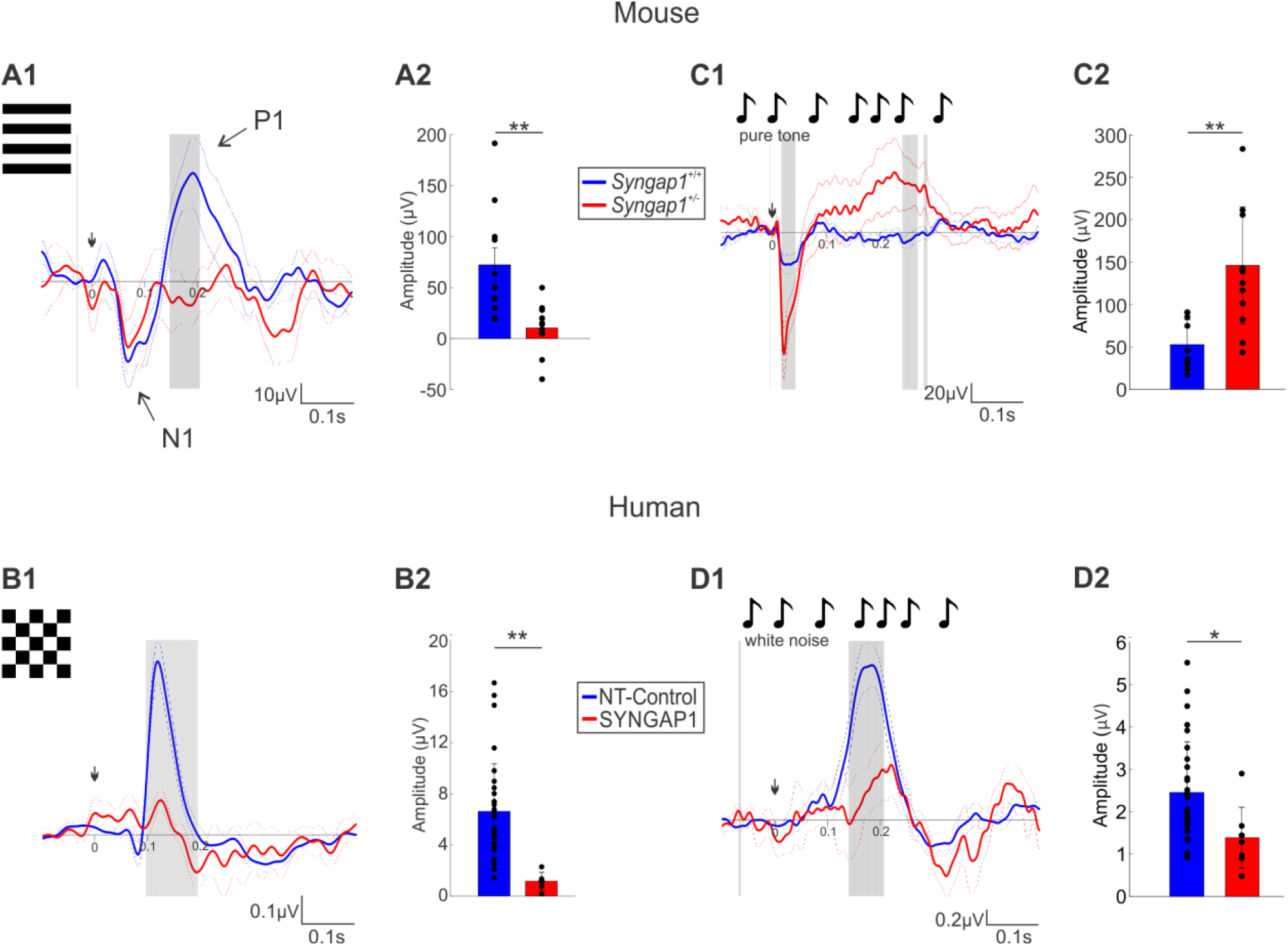
*Syngap1* haploinsufficiency correlates with decreased responses to visual stimuli in both patients and mice, while alterations of auditory responses are species-specific. **A1-B2**: VEP responses in mice (A1, A2) and humans (B1, B2). **A1, B1:** Grand average traces. Arrows show the start of stimulation. Shadow gray bars represent significant differences (p<0.05) between groups **A2, B2**: Bar plots shows P1 amplitude in (A2) mice (Wilcoxon rank sum test; z=-3.1884, p=0.0014) and (B2) humans (Wilcoxon rank sum test; z=3.7430, p=0.0001). **C1-D2**: AEP response in mice (C1, C2) and humans (D1, D2). *Syngap1^+/+^* N=11; *Syngap1^+/-^* n=11 mice, and NT Controls n=26; *SYNGAP1*-ID n=7 human participants. **C1, D1:** Grand average traces. **C2:** Bar plot shows N1 amplitude in mice (Wilcoxon rank sum test; z=-3.0863, p=0.0020). **D2**: Bar plot shows P1 amplitude in humans (Wilcoxon rank sum test; z=1.9899, p=0.0474). Bar graphs represent mean±SEM. *Syngap1^+/+^* n=11 and *Syngap1^+/-^* n=11 mice; NT Controls n=29 and *SYNGAP1*-ID n=8.

Next, we performed the same analyses on EEGs recorded from *SYNGAP1*-ID patients. Consistent with what we found in mutant mice (Figure 3A), we observed reduced P1 amplitude in VEPs recorded from *SYNGAP1*-ID patients as compared to the corresponding control group (Figure 3B). In contrast, AEP analysis highlighted different alterations within *Syngap1^+/-^* mice and *SYNGAP1*-ID patients. *SYNGAP1*-ID patients exhibit reduced P1, with no change in N1 (Figure 3D), while *Syngap1^+/-^* mice showed the opposite (compare Figure 3C and 3D).

Our results indicate that abnormalities in sensory-evoked related potentials do not provide a translationally robust biomarker for *SYNGAP1/Syngap1* haploinsufficiency, since they may be species-specific as well as dependent on the specific sensory modality.

Since the analysis of sensory-evoked local field potential amplitudes or delays did not show consistent differences across species (human and mice) and sensory perception modalities (auditory and visual), we decided to perform more sophisticated analyses. First, we asked whether there could be genotype-dependent effects on the temporal changes in distinct oscillation power over the frequency domain. Time frequency spectrograms (TFS) showed significantly different patterns between wild-type/neurotypic and *Syngap1/SYNGAP1* individuals (Figure 4, A1-D1). Although we could not identify a motif common to both mice and humans in any of the tested sensory modalities, we noted that gamma power seemed to be temporally organized by a lower rhythm in the theta band. To further investigate this possibility, we performed phase amplitude coupling (PAC) analysis. We found significantly increased PAC between theta and gamma bands, in both *Syngap1^+/-^* mice and *SYNGAP1*-ID patients, which was further consistent across sensory modalities (Figure 4A2-D2), suggesting that sensory stimulations lead to excessively coordination of the temporal organization of oscillatory activity in individuals carrying *Syngap1/SYNGAP1* pathogenic variants. Altogether, these results demonstrate that increased theta/gamma PAC evoked by sensory stimulation is a strong, reproducible biomarker of *Syngap1/SYNGAP1* haploinsufficiency.

**Figure 4.**
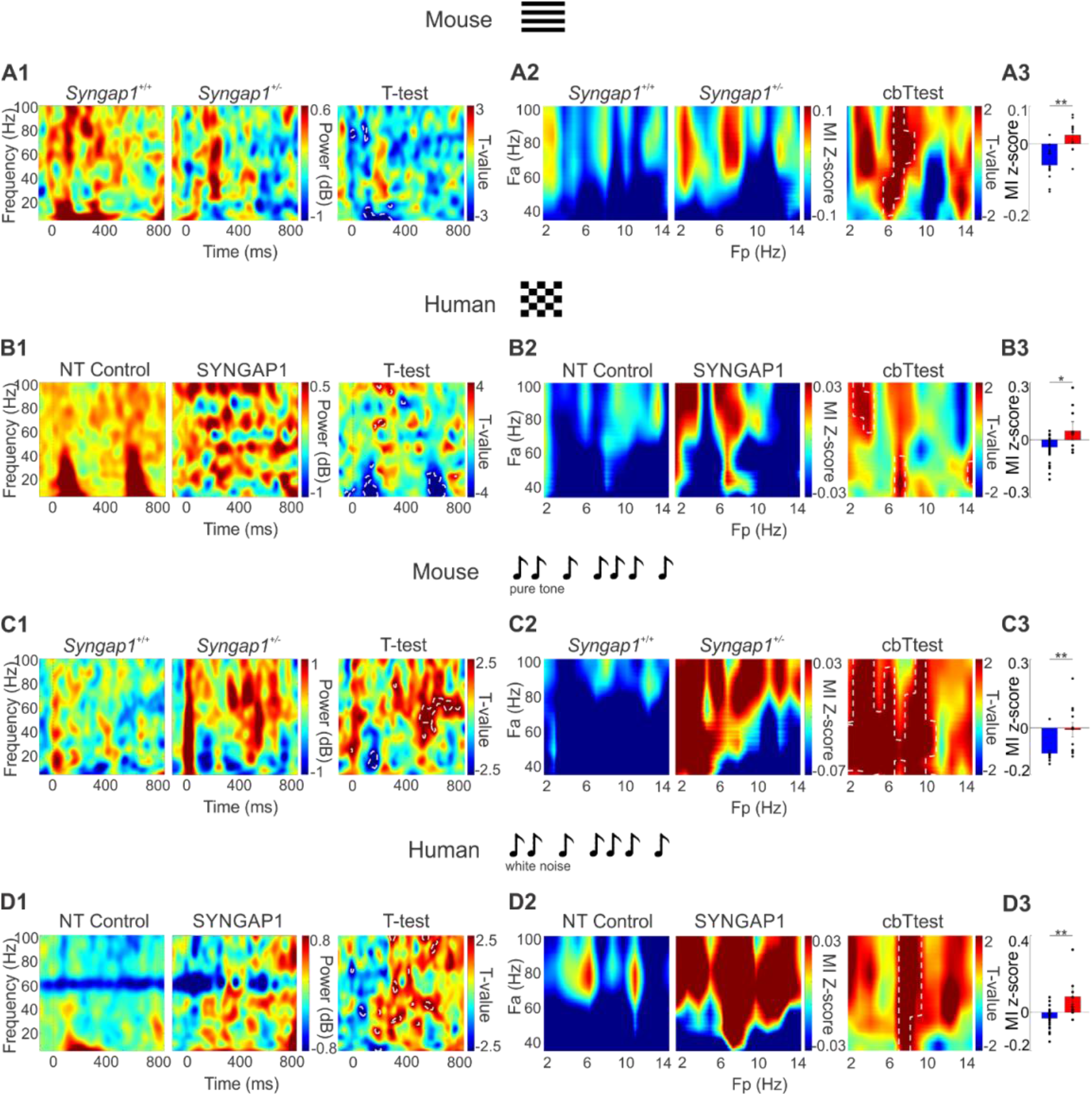
Theta/gamma phase-amplitude coupling (PAC) following auditory and visual stimuli is significantly increased in both *Syngap1^+/-^* mice and *SYNGAP1*-ID patients. **A1-D1**: Time Frequency Spectrogram showing visual (**A1, B1**) and auditory (**C1, D1**) evoked responses in mice (**A1, C1**) and humans (**B1, D1**). Dotted lines represent statistical differences (p<0.05) in Welch t-test map (right panels). Note that gamma power seems to be temporally organized by a lower frequency rhythm, which is particularly clear in B1. **A2-D2**: Phase-amplitude comodulograms and cluster based statistics. Statistical differences (p<0.05) are marked by white dotted lines. **A3-D3,** Bar plots shows z-score values corresponding to theta/gamma PAC. **A3**: t-test, T(20)=-4.1000, p=0.0005; **B3**: t-test, T(20)=-3.1898, p=0.0033; **C3**: Wilcoxon rank sum test; z=-3.0863, p=0.0020; **D3**: Wilcoxon rank sum test, z=-2.8813, p=0.0092; ** indicates p<0.01, * indicates p<0.05. A, C *: Syngap1^+/+^* n=11; *Syngap1^+/-^* n=11 mice; B: number of human participants, NT Controls n=26; *SYNGAP1*-ID n=7; D: : number of human participants, NT Controls n=29; *SYNGAP1*-ID n=8.

### Neural entrainment abnormalities are sensory-modality dependent in *Syngap1^+/-^* mice

When presented with rhythmic sensory stimulations oscillating at a certain frequency, neural networks oscillate in time to the stimulus, a process referred to as entrainment. Neural entrainment is thought to increase the ratio of signal to noise in the local cortical network, thus increasing sensory gain for a specific frequency^55^. Abnormal auditory entrainment following auditory stimulations has been reported in idiopathic ASD, and both in Fragile X syndrome patients^56–59^ and mouse models^25^. To explore whether similar neural entrainment abnormalities are caused by *Syngap1* haploinsufficiency, we recorded EEGs following repetitive auditory and visual entrainment protocols presented at different frequencies. Inter-trial coherence analysis showed significantly increased auditory entrainment following stimulation with click trains at 40Hz (Figure 5B), but not at 10Hz (Figure 5A), in *Syngap1^+/-^* mice compared to control littermates. In contrast, visual entrainment to stimuli (flickering lights) presented at 8 Hz (Figure 5C) was significantly reduced in mutant mice. However, visual entrainment following stimulation at 15Hz (Figure 5D) or 30 Hz (data not shown) was indistinguishable in *Syngap1^+/-^* versus control mice. No sex differences were identified in any of the entrainment paradigms (Supplementary Figure 5).

**Figure 5.**
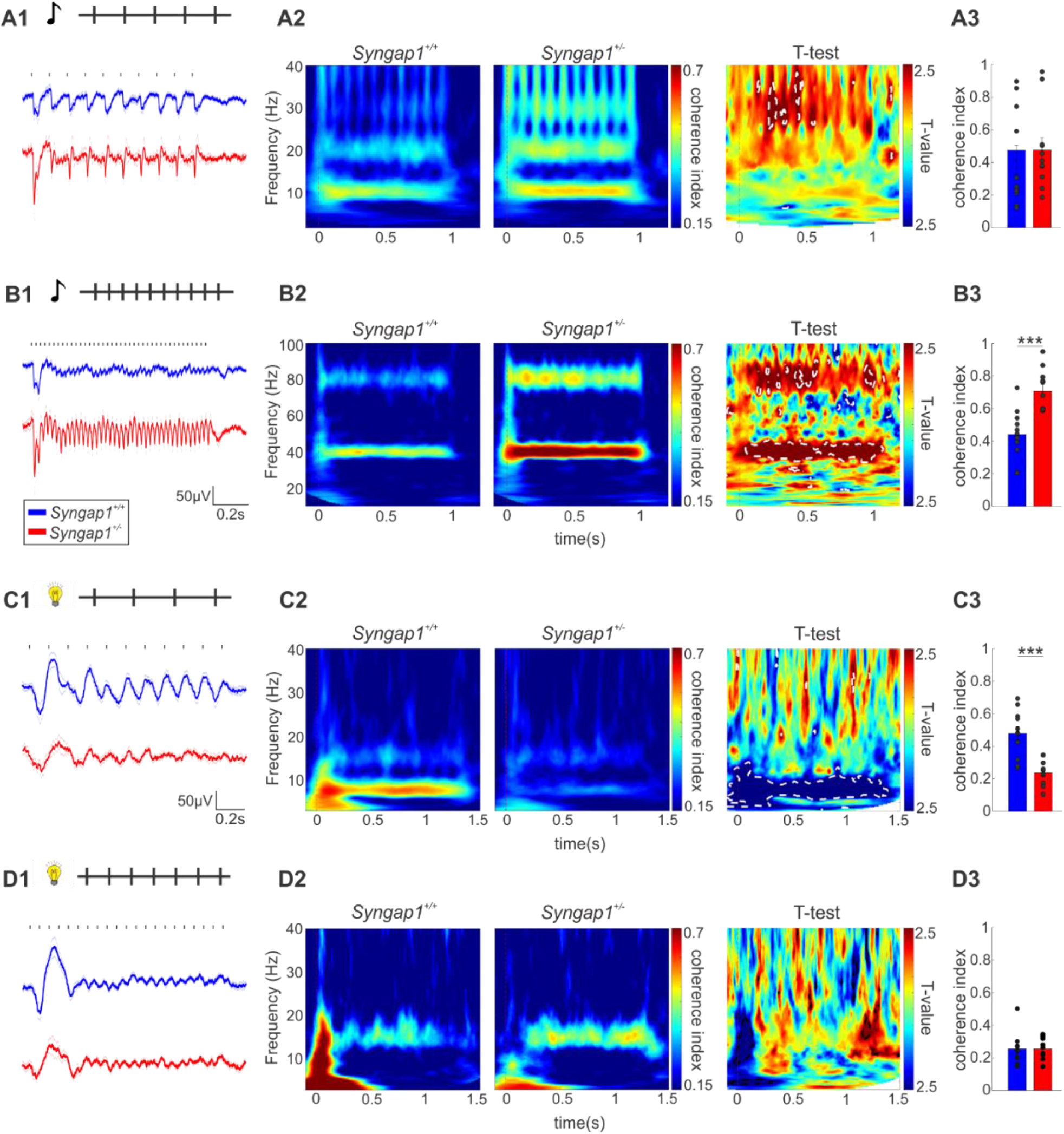
Syngap1 haploinsufficiency in mice is associated with excessive auditory entrainment at 40Hz but not at 10 Hz and reduced visual entrainment at 8Hz but not 15Hz. **A1-D1**: Grand average traces. On top, diagram showing the stimulation protocol: Auditory entrainment at 10 Hz (A1), at 40 Hz (B1) and visual entrainment at 8 Hz (C1) and 15Hz (D1). **A2-D2:** Inter-trial coherence and Welch t-test maps. Statistical differences (p<0.05) are marked by white dotted lines. **A3-D3**: Bar plots shows coherence index values corresponding to the stimulating frequency. **A3**: T-test, T(20)=0.5430, p=0.5931; **B3**: Wilcoxon rank sum test, z=-3.5459, p=0.0003; **C3**: Wilcoxon rank sum test; z=-3.4146, p=0.0006; **D3**: Wilcoxon rank sum test; z=-0.3583, p=0.7427 *** indicates p<0.001. *Syngap1^+/+^* n=11; *Syngap1^+/-^* n=11 mice.

These results suggest that neural entrainment alterations may be specific not only to particular frequencies but also to sensory modalities. The EEG recordings from *SYNGAP1*-ID patients that we had access to did not include stimulation protocols aimed at eliciting neural entrainment; however, our results suggest neural entrainment could be a potentially interesting biomarker to be explored in these patients.

Finally, we evaluated the synchronicity of brain oscillations to a chirp-modulated tone, since this stimulus has been reported to be reduced in Fragile X syndrome patients and mouse models^25, 57, 58^. Fragile X syndrome is a neurodevelopmental disorder, which could potentially share some pathophysiological mechanisms with the *Syngap1* haploinsufficiency^60^. We tested both up and down chirps to ensure that the differences are specific to modulation frequency and not affected by the direction of frequency change in the sound. In contrast to previous observations in Fragile X Syndrome, *Syngap1^+/-^* male mice showed an increased response to up (Supplemental Figure 6) and down (data not shown) chirp stimulation. Strikingly, in our experiments, neural responses to a chirp stimulus were sex dependent, since we could not induce neural entrainment with a chirp stimulus neither in wild-type nor in mutant female mice (Supplemental Figure 6). Given the lack of reproducible response to a chirp stimulus in females as compared to males, this particular stimulation paradigm may not be an appropriate translational biomarker.

### *Syngap1/SYNGAP1* haploinsufficiency in associated with decreased habituation to repetitive auditory stimuli

Habituation responses to repeating tones (auditory gating) or novelty detection responses (mismatch negativity) have been largely employed as biomarkers in neurodevelopmental disorders^61^. Therefore, we investigated whether *Syngap1^+/-^* mice show deficits in habituation, by analyzing cortical response to 5-9 consecutive sounds presented at a fixed repetition rate of 1Hz (Figure 6A1). In particular, we measured N1 amplitude of AEP evoked by the first (S1) and the second sound (S2). In contrast to the clear reduction of N1 amplitude observed in *Syngap1^+/+^* mice, *Syngap1^+/-^* mice showed no change in N1 amplitude between S1 and S2 (Figure 6 A2, A3), and this phenotype was similar in both sexes (Supplementary Figure 7).

**Figure 6.**
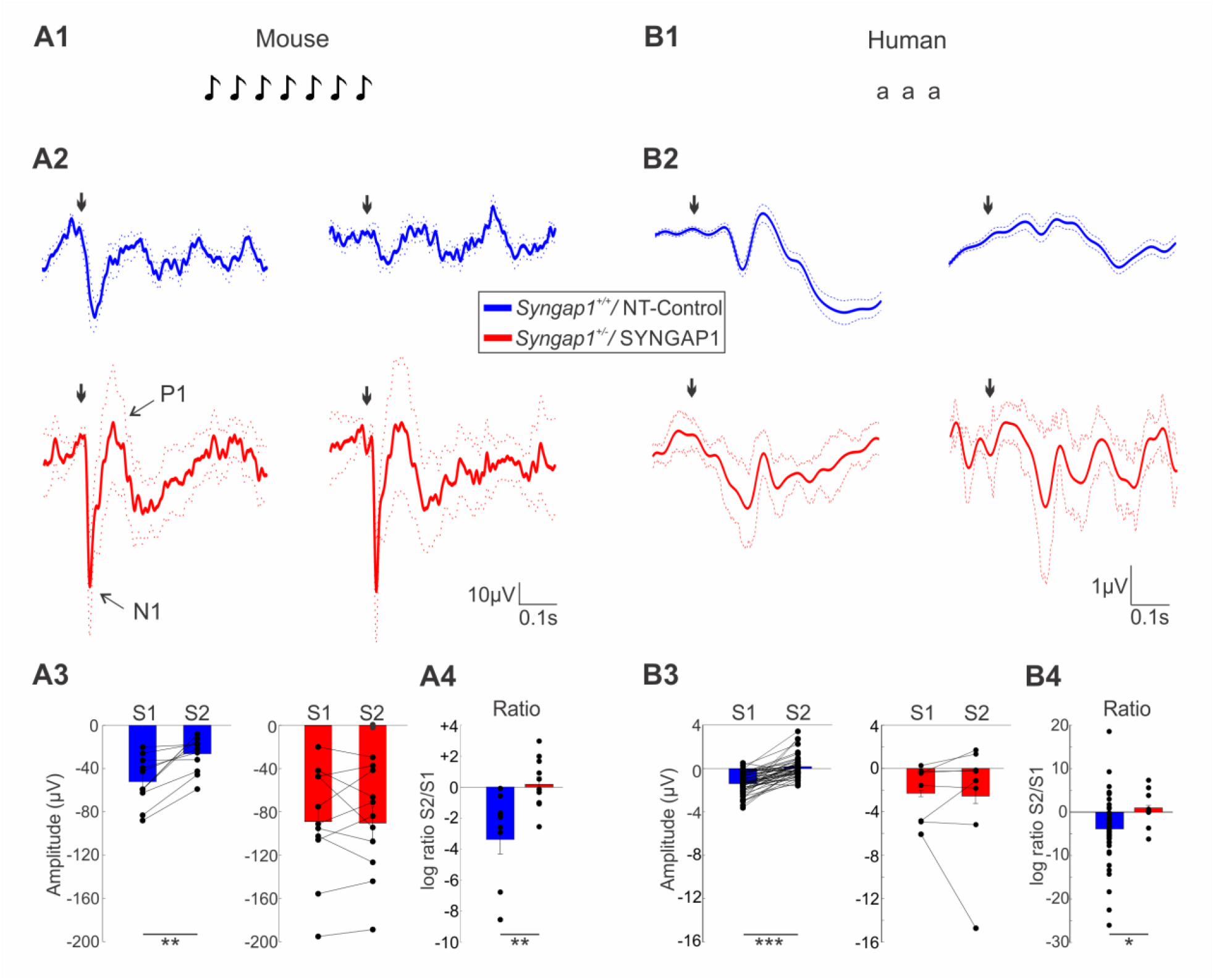
*Syngap1/SYNGAP1* haploinsufficiency is associated with a lack of habituation to repetitive sounds in both mice and humans. **A1-B2:** Schematic of repetitive auditory stimulation protocol in mice (A1) and humans (B1). **A2, B2:** AEP response to the first (left) and second (right) standard sound of *Syngap1^+/+^* (blue) vs *Syngap1^+/-^* (red) mice (A1) and NT-Controls vs *SYNGAP1*-ID individuals (B1). **A3, B3:** Bar plots show P1 amplitude after first (S1) vs second (S2) sounds in mice (A2; Syngap1^+/+^ mice, Wilcoxon signed rank, z=-2.9355, p=0.0033 and *Syngap1^+/-^* mice, Wilcoxon signed rank, z=0.1778, p=0.8589) and humans (B2; NT-Control individuals, paired t-test, T(48)=-7.6659, p<0.0001 and *SYNGAP1*-ID individuals, paired t-test, T(7)=0.12209, p=0.9141). **A4, B4:** Logarithmic ratio S2/S1 in mice (Wilcoxon rank sum test; z=-3.0214, p=0.0025) and humans (Welch independent t-test; T(14)=0.0282). *Syngap1^+/+^* n=11; *Syngap1^+/-^* n=11 mice; number of human participants, NT Controls n=49; *SYNGAP1*-ID n=8.

In keeping with our observation in the mouse model, we observed a similar lack of habituation in *SYNGAP1*-ID individuals (Figure 6B2, B3). Taking into account the difference between absolute N1 amplitudes between different mice or individuals, we calculated the S2/S1 ratio for each subject (Figure A4, B4) and found a consistent lack of habituation in both *Syngap1^+/-^* mice and *SYNGAP1*-ID patients, suggesting that it could be a robust translational biomarker for *Syngap1/SYNGAP1* haploinsufficiency

Mismatch negativity (MMN) is a memory-based brain response to any discriminable change in a stream of auditory stimulation^62^. MMN has recently gained attention as a tool for neurological evaluations since several studies indicate that it is a robust marker of cognitive dysfunction^63^. Here, we explored whether the ability to detect actual physical difference in sounds was altered in *Syngap1^+/-^* mice, by using a sequence of repetitive, standard sensory stimuli, interrupted by an oddball or deviant stimulus (Figure 7A). We found that the mean amplitude of the first negative peak of MMN (N1), which indicates the sound-specific responses to the auditory stimuli presented, was not statistical different between genotypes (Figure 7C, D). Conversely, while the MMN pattern in wild-type mice showed the expected positive peak at 70-110 ms after the stimulus presentation (P1), indicating that the deviant sound was detected correctly, this peak was wholly absent in *Syngap1^+/-^* mice (Figure 7C, D). No sex differences were observed for the described MMN pattern (Supplementary Figure 7). Overall, these results suggest that, while responses to frequence specificity are intact at the primary sensory level, deviant sound detection, known to rely on higher-order sensory functions, is specifically altered in *Syngap1^+/-^* mice.

**Figure 7.**
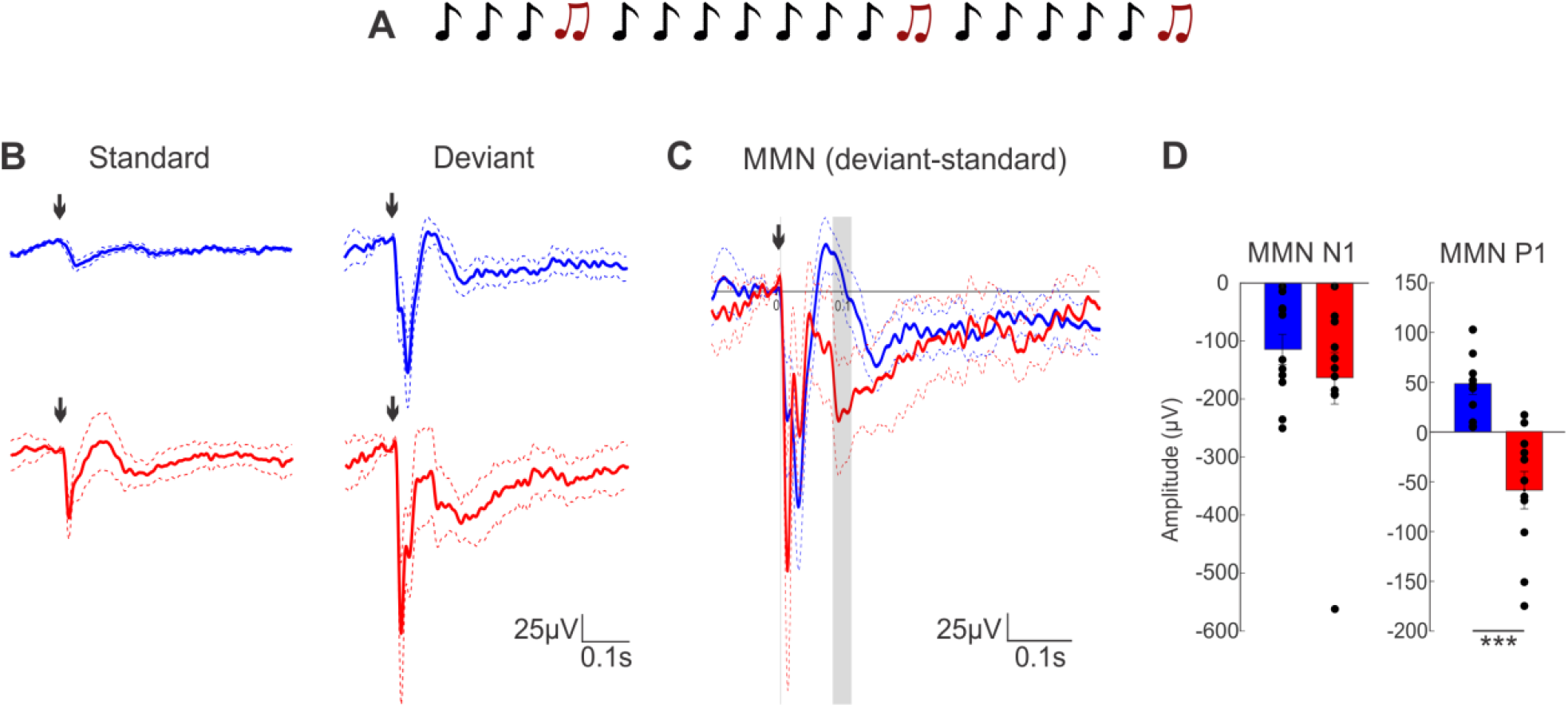
*Syngap1* haploinsufficiency is associated with an abnormal MMN pattern in mice. **A:** Schematic of Oddball stimulation protocol in mice. **B:** AEP traces of the standard (left) and deviant (right) sounds of *Syngap1^+/+^* (blue) and *Syngap1^+/-^* (red) mice. **C:** superposed MMN (deviant - standard) traces. **D:** Bar plots of MMN N1 (Wilcoxon rank sum test; z=-0.7223, p=0.4701) and MMN P1 (Wilcoxon rank sum test; z=-3.5469, p=0.0003) peak amplitude. Shadow gray bars represent significant differences (p<0.05) between groups. Arrows indicate the start time of stimulation. ** indicates p<0.01, *** indicates p<0.001. *Syngap1^+/+^* n=11; *Syngap1^+/-^* n=11 mice.

## Discussion

*Syngap1* haploinsufficiency is associated with intellectual disabilities, developmental delay, epilepsy, poor or absent language development, autistic traits and other behavioral alterations. Sensory processing and perception have been recently shown to contribute to cognitive and behavioral dysfunction in specific neurodevelopmental disorders, including ASD; however, whether and to what extent visual and auditory sensory processing is affected in *SYNGAP1*-ID or in related mouse models remained unclear. Here, we identified, in *Syngap1^+/-^* mice, specific alterations in multiple aspects of auditory and visual processing, including increased baseline gamma oscillation power, increased theta/gamma phase amplitude coupling following stimulus presentation and lack of habituation to repetitive auditory stimuli. These alterations were also present in the human *SYNGAP1*-ID population as well, suggesting that they may be valid as reliable translational biomarkers of sensory processing alterations associated with *SYNGAP1/Syngap1* haploinsufficiency. In the *Syngap1^+/-^* mouse model, we further found increased interictal activity during immobility, abnormal neural entrainment at different, sensory modality-specific frequencies and abnormal deviant sound detection. These phenomena could be explored in the human patient population as well to determine to what extent these alterations are conserved.

### Interictal activity during immobility

One of the most frequent neurological problems in *SYNGAP1*-ID patients is the occurrence of epileptic encephalopathy. Extensive work showed the presence of epileptic phenotypes in different mouse models of *Syngap1* haploinsufficiency^14, 15, 43^. Classically, researchers have focused on seizure duration, and their frequency, however seizures may appear with a low frequency and their observation requires either long-term recording^15^ or the need to induce them using drugs or sensory stimulation^14^. Interictal spikes are pathological electrical events that reflect seizure susceptibility in patients and share common mechanisms with seizures^44^. In the *Syngap1^+/-^* mouse model, we found a relatively high frequency of interictal spikes, which was already evident in the first 15 minutes of recording for all *Syngap1^+/-^* mice and in the different brain regions examined. This finding is consistent with what is previously reported in two different *Syngap1* haploinsufficient mice models as well as in *SYNGAP1*-ID patients^14, 15^. Therefore, inter-ictal events, in particular during sleep and behavioral state transitions, could be a reliable and easy to measure biomarker for the evaluation of the efficacy of drug treatments on cortical hyper-excitability in *SYNGAP1* associated disorders.

### Translational biomarkers in baseline activity and sensory processing

Event Related Potentials (ERP) are frequently used as quantitative methods to directly compare sensory processing in patients with neurodevelopmental disorders and the corresponding mouse models^16, 17^, nevertheless we did not find a consistent ERP phenotype either across sensory modalities or between species in the case of *Syngap1/SYNGAP1* haploinsufficiency. A potential confounding effect is the use of different anti-epileptic medications in *SYNGAP1*-ID patients, which could affect ERP dynamics or amplitude. On the other hand, PAC analysis performed in the same data set revealed significant and consistently increased theta/gamma PAC following visual and auditory stimulation in *Syngap1^+/-^* mice and *SYNGAP1*-ID patients. Whether the augmented PAC is triggered by sensory stimulation in *Syngap1^+/-^* mice and *SYNGAP1*-ID patients or whether it reflects a basal disorganised network remains to be elucidated. Nevertheless, while PAC is an emerging tool mostly used to study local and long range connectivity^64^, recent studies support its potential interest as an EEG biomarker of different neurological pathologies^41, 65, 66^. For example, enhanced PAC has been extensively reported in epileptic patients and mouse models of epilepsy^65, 67–69^, thus leading to the hypothesis that increased PAC reflects network hyper-excitability^70^. Increased PAC has also been reported in other disorders such as obsessive-compulsive disorder^71^ and Parkinsons disease^72^. In particular, PAC levels appear proportional to the severity of motor symptoms in Parkinsons disease^66, 72^, suggesting that excessive PAC “may reflect a pathological state in which the cortex is restricted to a monotonous pattern of coupling, rendering it less able to respond dynamically to signals from other cortical regions, such as frontal executive areas involved in internally directed movement”^72^. A similar rigid network organization could underlie the increased PAC observed in *Syngap1/SYNGAP1* haploinsufficiency.

Here, we observed a consistent increase in baseline gamma band power across all the regions we recorded in *Syngap1* mutant mice and *SYNGAP1*-ID patients. A recent study found alterations in the gamma power modulation during behavioral-state transitions in *Syngap1^+/−^* mice^15^. Interestingly, this same study revealed a higher occurrence of myoclonic seizures during behavioral-state transitions in *Syngap1^+/-^* mice and possibly *SYNGAP1*-ID patients^15^. Altogether, these observations highlight a potential link between the two phenomena, which could share common pathophysiological mechanisms. Indeed, several studies have shown that networks characterized by more prominent gamma oscillations are more likely to generate ictal activity, thus suggesting that increased gamma power may play a role in seizure generation^73, 74^. For instance, in photosensitive epileptic patients, an increase in phase synchrony in the gamma-band (30-120 Hz) preceded an increased synchronization during flicker stimulation, but only when such stimulation lead to epileptic photoparoxysmal responses^75^. Of note, two recent studies showed that <70% of awake seizures occurred at eye closure or with eyes closed in a cohort of *SYNGAP1*-ID patients with developmental epileptic encephalopathy^76, 77^, supporting the hypothesis that cortical networks may be more prone to seizures even in absence of stimulation in *SYNGAP1*-ID.

Enhanced baseline gamma power has been described also in the auditory cortex of Fragile X syndrome and related mouse models (*Fmr1* KO mice)^25, 58, 78^. Using an auditory chirp stimulation, the same studies showed that inter-trial phase synchrony was reduced in both humans with Fragile X syndrome and *Fmr1* KO mice, particularly at gamma frequency^25, 58^. These data lead to the hypothesis that enhanced background gamma oscillation, or network noise, may interfere with stimulus-evoked synchronization in Fragile X syndrome. In contrast, *Syngap1^+/−^* mice showed enhanced inter-trial phase synchrony when challenged with either a chirp or rhythmic auditory stimulation at 40Hz, but not 10Hz, suggesting that increased baseline gamma band power does not directly lead to deficits in mounting a specific gamma frequency-locked response to sounds. On the other hand, we observed significantly reduced entrainment in the visual cortex of *Syngap1^+/−^* mice specifically following a rhythmic sensory stimulation at 8Hz. These observations suggest that different neural circuit changes may underlie distinct neural entrainment alteration in visual versus auditory cortex. Neural entrainment may enhance signal-to-noise ratio in the local cortical network, thus increasing sensory gain for the specific frequency^55^. Thus, different region-specific neural entrainment alterations (enhanced in auditory cortex versus weakened in visual cortex) in *Syngap1^+/-^* mice may indicate different effects on sensory processing and perception depending on the sensory modality.

In line with this hypothesis, we observed increased AEP amplitude responses to pure tones in auditory cortex, but reduced VEP amplitude responses to visual stimulation in *Syngap1^+/−^* mice. In addition, a recent study showed hyposensitivity to touch in somatosensory cortex in another *Syngap1^+/−^* mutant model^22^. These observations suggest that *Syngap1* haploinsufficiency does not lead to generalized hyper or hyposensitivity to sensory stimuli, but to cortical region-specific deficits, possibly relying on diverse distinct neuronal circuit alterations.

### Potential contribution of GABAergic circuit deficits to sensory processing alterations

While the cellular bases underlying the generation of specific brain rhythms are not yet completely understood, extensive work point to a critical role of GABAergic interneurons, in particular parvalbumin-expressing (PV) cells, the larger subgroup of cortical GABAergic cells, which target the perisomatic domain of pyramidal neurons, and somatostatin (SST)-expressing, dendritic targeting interneurons^79–81^. SYNGAP1 protein is expressed by GABAergic cells and *Syngap1* deletion in single PV cells in cortical organotypic cultures lead to reduced perisomatic synapse density^82^. Several of the specific EEG alterations we observed in *Syngap1^+/-^* mice are consistent with PV cell circuit deficits. First, PV cells play a key role in generating and maintaining gamma oscillations in the brain ^83, 84^. In particular, recent studies have also shown that deficiency in PV interneuron-mediated inhibition contribute to increased baseline cortical gamma rhythm^79, 85, 86^, a phenotype we observed in both mutant mice and *SYNGAP1*-ID patients (Figure 2). Second, auditory habituation has been shown to involve an increase in inhibition^87, 88^. Specifically, PV cells have been shown to amplify habituation providing non-specific inhibition. For instance, optogenetic suppression of PV cells led to increased responses to both repetitive (standard) and rare (deviant) tones^87^, which is similar to what we observed in *Syngap1^+/-^* mice (Figure 7). Third, recent studies showed that optogenetically suppressing PV cell activity increased theta/gamma coupling in visual cortex^79^, suggesting that increased theta/gamma PAC (Figure 4) might indicate reduced PV cell activity. Further studies will be needed to clarify how and to what extent cortical PV cell circuit connectivity and function are altered by *Syngap1* haploinsufficiency and whether these alterations might be different in different cortical regions.

Furthermore, we cannot exclude the involvement of different interneuron-specific dysfunctions in *Syngap1* haploinsufficiency. For example, SST interneurons selectively reduced excitatory responses to frequent tones in an oddball paradigm^87^ and optogenetic suppression of SOMs cells reduced the MMN peak^89^, suggesting impaired deviant sound detection, similarly to what we observed in *Syngap1^+/-^* mice (Figure 7). Altogether these observations point to the contribution of GABAergic interneurons to the observed sensory perception phenotypes in *Syngap1/SYNGAP1* haploinsufficiency. Further studies are needed to address the specific GABAergic network alterations and their relative contribution to different aspects of sensory processing and perception.

In conclusion, this study shows that visual and auditory perception is altered in *Syngap1/SYNGAP1* haploinsufficiency, and reveal multiple novel robust and translation relevant phenotypes, which could both be used as biomarkers to monitor therapeutic outcomes of treatments and provide opportunities to better understand the pathophysiology of *Syngap1/SYNGAP1* haploinsufficiency.

## ACKNOWLEDGEMENTS

We thank Dr. Jean Marc Lina for his insightful suggestions on PAC analysis. We would like to thank James Bellord Waldron and Antônia Samia Fernandes do Nascimento for their technical assistance, the Comité Institutionnel de Bonne Pratiques Animales en Recherche (CIBPAR), all the personnel of the animal facility of the Research Center of CHU Sainte-Justine (Université de Montreal) and the iPED-NeuroD human EEG platform (S.L.) for their instrumental technical support.

