## Supplemental Material for "Sensory processing dysregulations as reliable translational biomarkers in *SYNGAP1* haploinsufficiency"

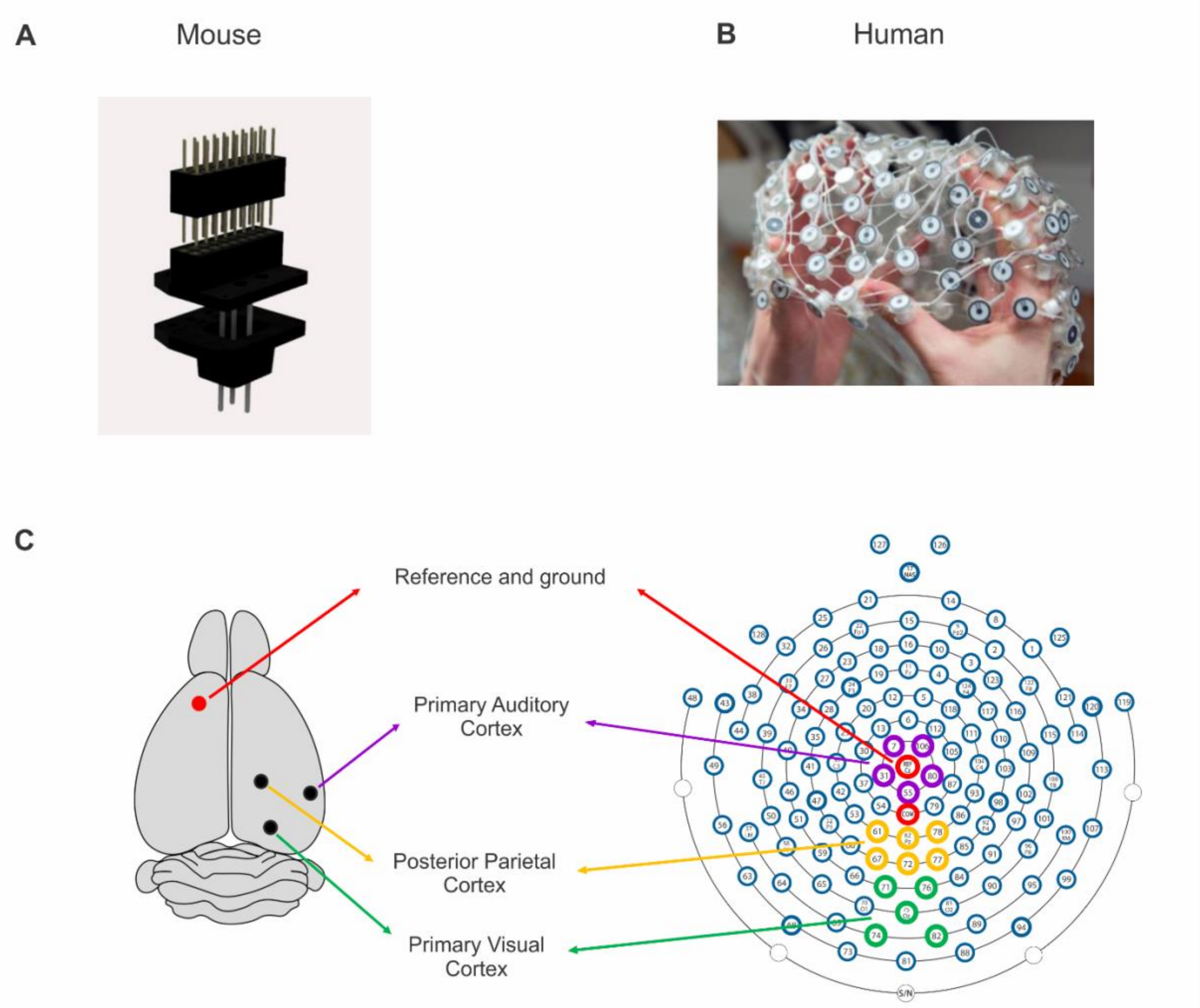


**Supplementary Figure 1. Electrodes used in mice and humans. A,** Scheme of the custom-designed 3D printed microdrive-like scaffold that holds the 3 electrode clusters at fixed positions. **B**, Geodesics 128 electrode nets used in humans. **C**, Schematic comparing the recording sites in mice (left panel) and human participants (right panels) analyzed in this study.

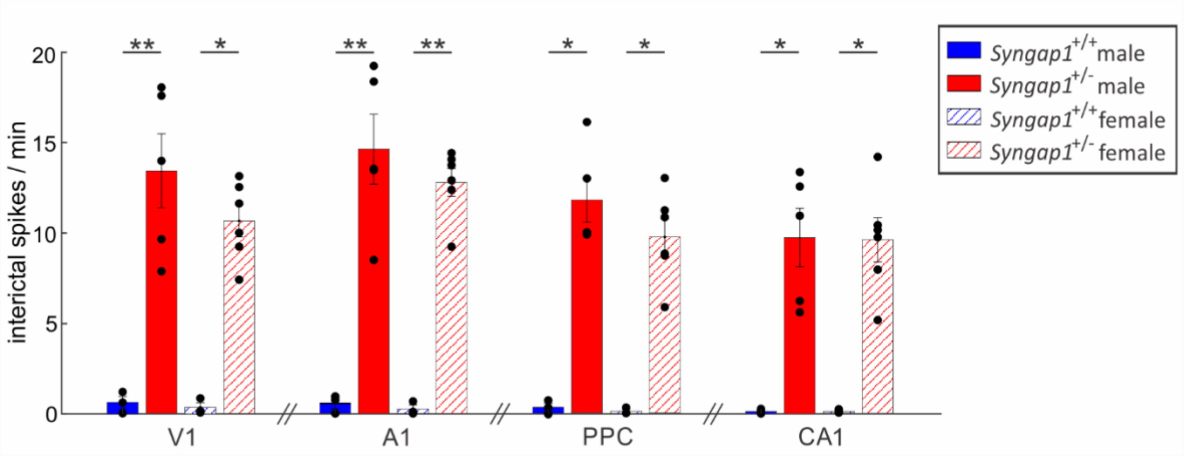


**Supplementary Figure 2. Inter-ictal spike activity is similar in male and female *Syngap1*$^{+/-}$ mice.** Bar plots show the total number of interictal spikes detected in different regions. **V1**: Two-way ANOVA; $F_{genotype}$ (1,13)=61.9908, p<0.0001, $F_{sex}$(1,13)=1.1014, p=0.3131, $F_{genotype*sex}$(1,13)=0.7136, p=0.4135. **A1**: Two-way ANOVA; $F_{genotype}$ (1,13)=94.4883, p<0.0001, $F_{sex}$(1,13)=0.6268, p=0.4427, $F_{genotype*sex}$(1,13)=0.3180, p=0.5824. **PPC**: Two-way ANOVA; $F_{genotype}$ (1,13)=91.5281, p<0.0001, $F_{sex}$(1,13)=1.1240, p=0.3084, $F_{genotype*sex}$(1,13)=0.6506, p=0.4344. **CA1:** Two-way ANOVA; $F_{genotype}$ (1,13)=48.0297, p<0.0001, $F_{sex}$(1,13)=0.0060, p=0.9397, $F_{genotype*sex}$(1,13)=0.00005, p=0.9818. Tukey *posthoc* test showed statistical differences between males and females in all recorded regions (**V1**: males p=0.0026, females p=0.0126; **A1**: males p=0.0010, females p=0.0026; **PPC**: males p=0.0110, females p=0.0272, **CA1**: males p=0.0139, females p=0.0126). * indicates p<0.05 and ** indicates p<0.01. Bar graphs represent mean±SEM. Mice numbers: Male *Syngap1*$^{+/+}$ n=3; Male *Syngap1*$^{+/-}$ n-=5; Female *Syngap1*$^{+/+}$ n=3; Female *Syngap1*$^{+/-}$ n=6.

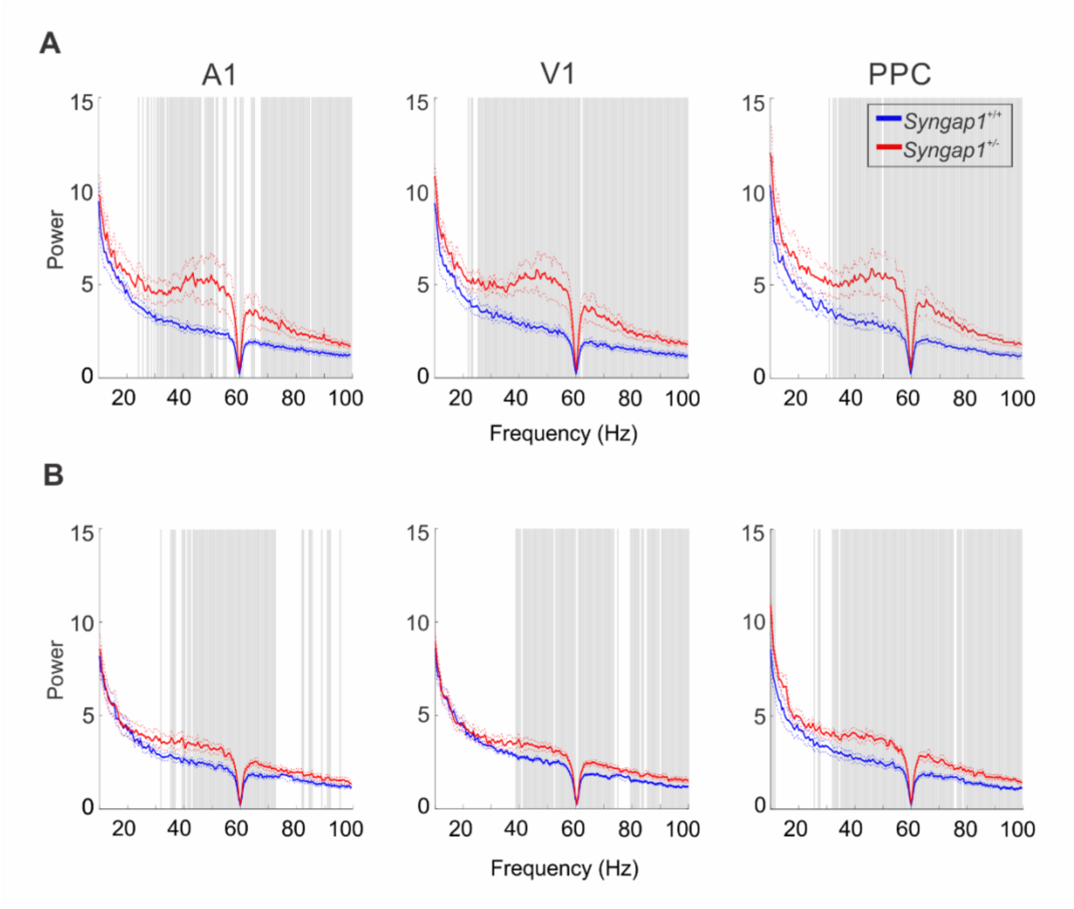


**Supplementary Figure 3. Increased baseline gamma power is present in both males and females *Syngap1*$^{+/-}$ mice. A-B**: Power Density Spectra of 3-minutes baseline state EEG signal from A1, V1 and PPC, in males (A) and females (B). Shadow gray bars represent statistical differences between groups (t-test, $p<0.05$). Mice numbers: Male *Syngap1*$^{+/+}$ n =5; Male *Syngap1*$^{+/-}$: n=5; Female *Syngap1*$^{+/+}$ n=6; Female *Syngap1*$^{+/-}$ n=6.

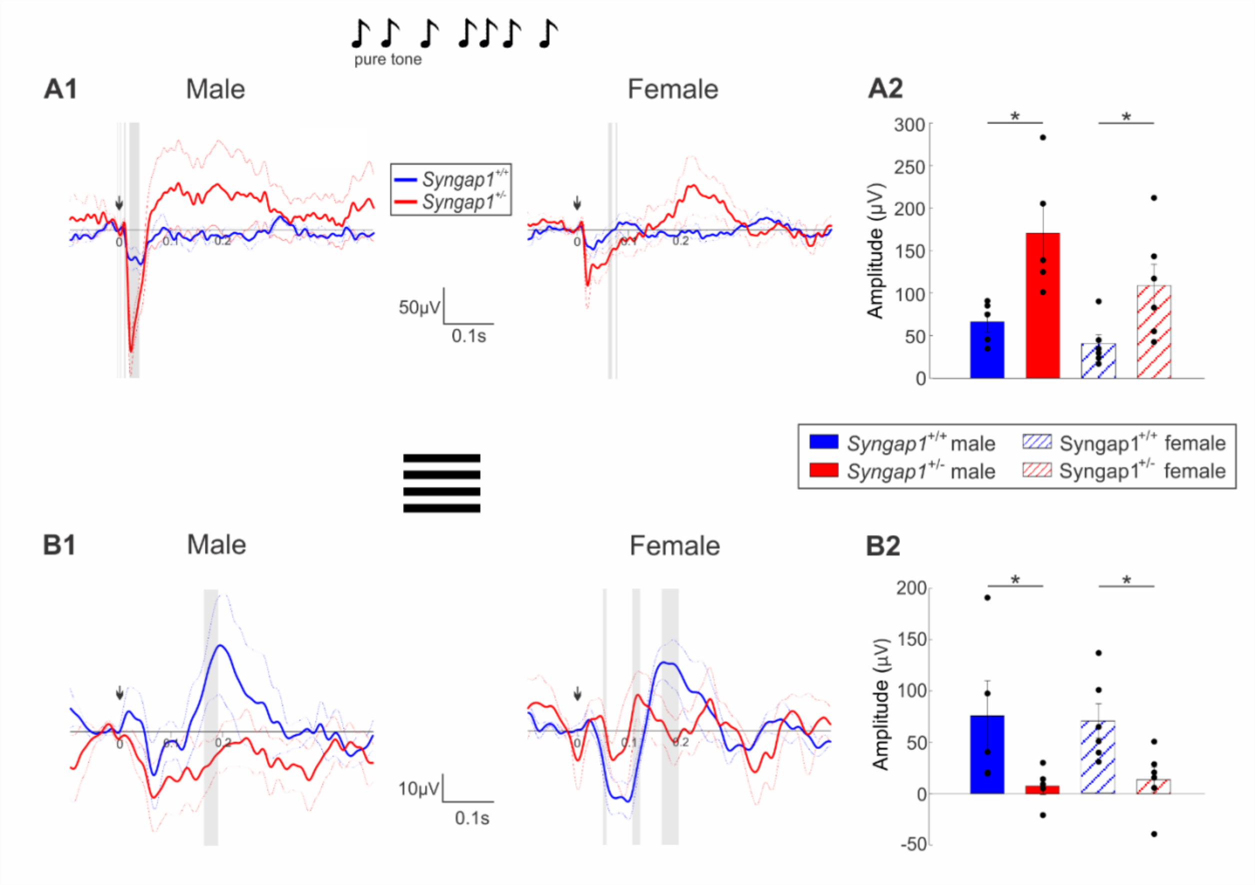


**Supplementary Figure 4. Visual and auditory responses do not show sex-related differences in mice.** **A1-B2**: AEP response. **A1, B1:** Grand average traces. Arrows show the start of stimulation. Gray bars represent significant differences ($p<0.05$) between groups **A2:** Bar plots shows N1 amplitude. Two-way ANOVA, $F_{genotype}$ (1,18)=15.3231, p=0.0010, $F_{sex}$(1,18)=3.9887, p=0.0612, $F_{genotype*sex}$(1,18)=0.7357, p=0.4023. Tukey *posthoc* test showed statistical differences in males (p=0.0046) and females (p=0.0360) **B2:** Bar plots shows P1 amplitude. Two-way ANOVA, $F_{genotype}$ (1,18)=10.1304, p=0.0052, $F_{sex}$(1,18)=0.00003, p=0.9953, $F_{genotype*sex}$(1,18)=0.0784, p=0.7826. Tukey *posthoc* test showed statistical differences in males (p=0.0307) and females (p=0.0452). Bar graphs represent mean±SEM. Mice numbers: Male *Syngap1*$^{+/+}$ n =5; Male *Syngap1*$^{+/-}$: n=5; Female *Syngap1*$^{+/+}$ n=6; Female *Syngap1*$^{+/-}$ n=6.

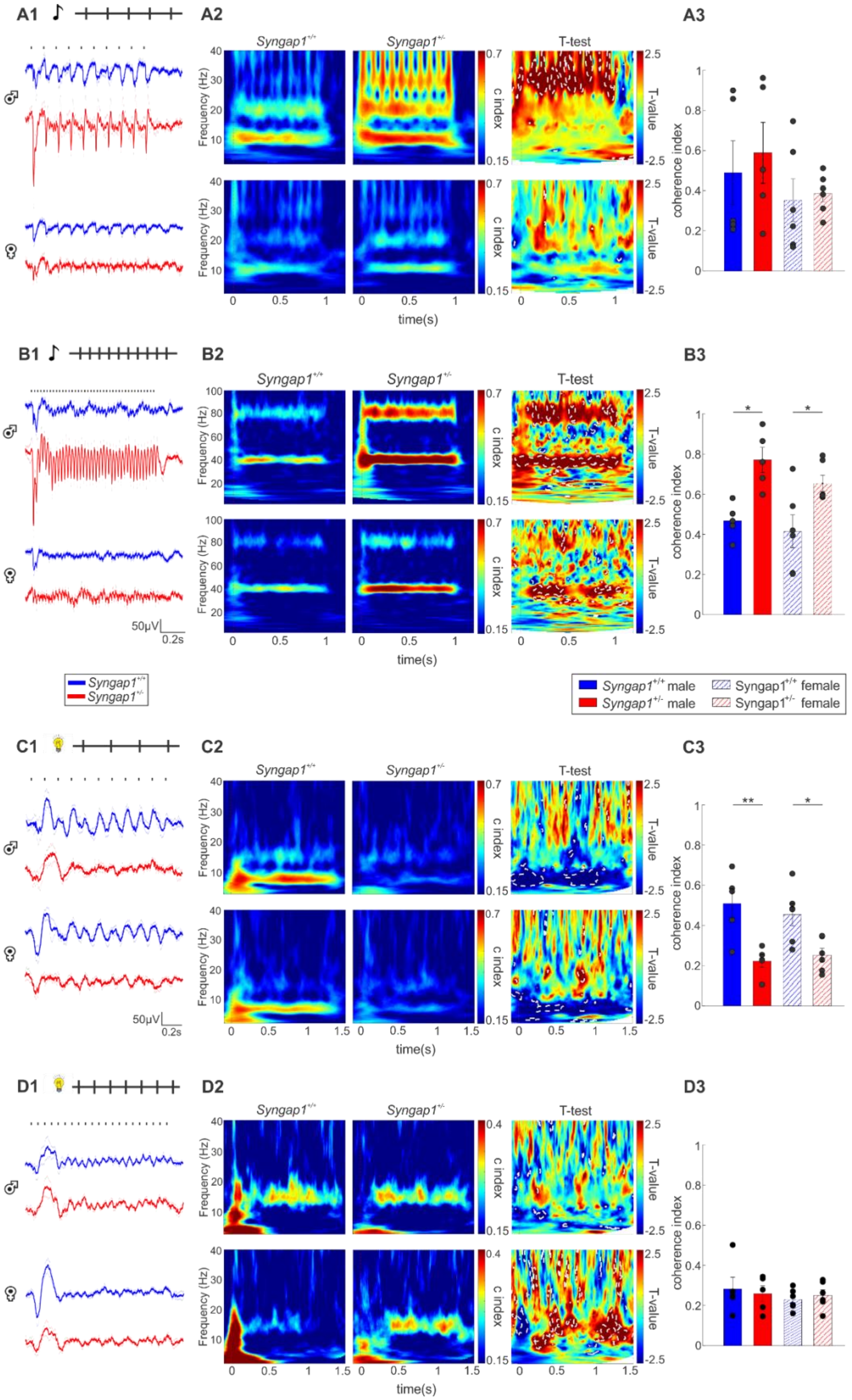

A1
A2
A3
B1
B2
B3
C1
C2
C3
D1
D2
D3
Syngap1+/+
Syngap1+/-
T-test
Frequency (Hz)
c index
T-value
time(s)
coherence index
50μV
0.2s
Syngap1+/+ male
Syngap1+/- male
Syngap1+/+ female
Syngap1+/- female

**Supplementary Figure 5. Visual and auditory entrainment alterations are comparable in *Syngap1*$^{+/-}$ male and female mice. A1-D1**: Grand average traces. On top, diagram showing the stimulation protocol: auditory entrainment at 10 Hz (A1), at 40 Hz (B1) and visual entrainment at 8 Hz (C1) and 15Hz (D1). Lines on top of traces represent each click of the train. **A2-D2:** Inter-trial coherence and Welch t-test maps. Statistical differences ($p<0.05$) are marked by white dotted lines. **A3-D3**: Bar plots shows coherence index values corresponding to the stimulating frequency. **A3:** Two-way ANOVA, $F_{genotype}$ (1,18)=0.3239, p=0.5763, $F_{sex}$(1,18)=1.8895, p=0.1861, $F_{genotype*sex}$(1,18)=0.0793, p=0.7816 and Tukey *posthoc* test for main effect in males (p=0.9380) and females (p=0.9964); **B3**: Two-way ANOVA, $F_{genotype}$ (1,18)=21.4249, p=0.0002, $F_{sex}$(1,18)=1.7241, p=0.2057, $F_{genotype*sex}$(1,18)=0.4541, p=0.5090 and Tukey *posthoc* test for main effect in males (p=0.0103) and females (p=0.0404); **C3:** Two-way ANOVA, $F_{genotype}$ (1,18)=22.9721, p=0.0001, $F_{sex}$(1,18)=0.0700, p=0.7943, $F_{genotype*sex}$(1,18)=0.6436, p=0.4329 and Tukey *posthoc* test for main effect in males (p=0.0067) and females (p=0.0383); **D3:** Two-way ANOVA, $F_{genotype}$ (1,18)=0.0015, p=0.9691, $F_{sex}$(1,18)=0.6683, p=0.4243, $F_{genotype*sex}$(1,18)=0.3224, p=0. 5772 and Tukey *posthoc* test for genotype effect in males (p=0.9759) and females (p=0.9789). Mice numbers: Male *Syngap1*$^{+/+}$ n=5; Male *Syngap1*$^{+/-}$: n=5; Female *Syngap1*$^{+/+}$ n=6; Female *Syngap1*$^{+/-}$ n=6.

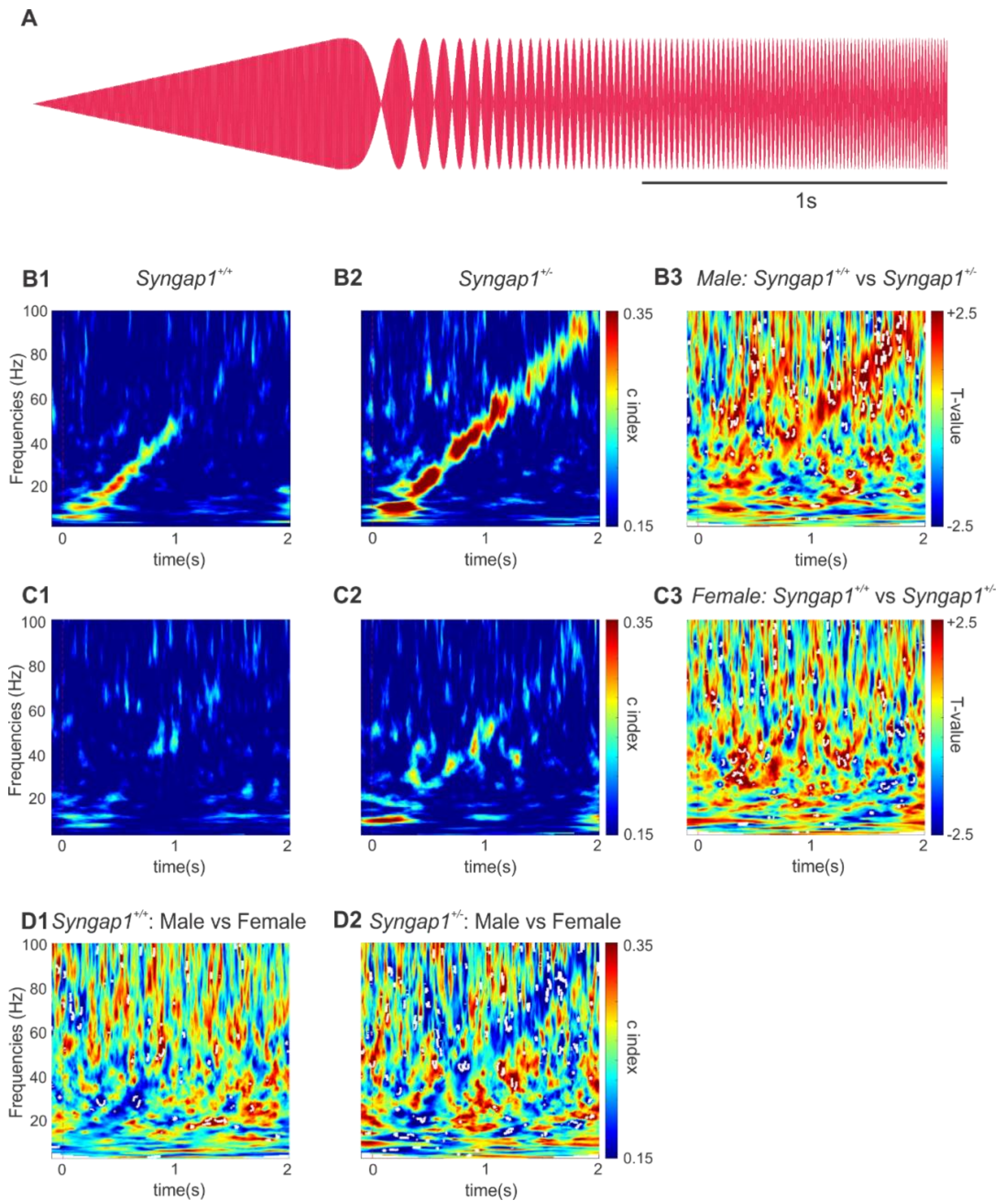


**Supplementary 6. Phase-locking response to chirp auditory stimuli is increased in Syngap1$^{+/-}$ male mice but is absent in females of either genotypes. A1**: Example of "Up Chirp" (0-100Hz) auditory stimuli. **B1, B2, C1, C2**: Inter-trial coherence maps of male *Syngap1*$^{+/+}$ (**B1**), male *Syngap1*$^{+/-}$ (**B2**), female *Syngap1*$^{+/+}$ (**C1**) and female *Syngap1*$^{+/-}$ mice (**C2**). **B3, C3, D1, D2**: Welch t-test maps comparing male *Syngap1*$^{+/+}$ vs male *Syngap1*$^{+/-}$ (**B3**), female *Syngap1*$^{+/+}$ vs female *Syngap1*$^{+/-}$ (**C3**), male *Syngap1*$^{+/+}$ vs female *Syngap1*$^{+/+}$ (**D1**) and male *Syngap1*$^{+/-}$ vs female *Syngap1*$^{+/-}$ (**D2**). Statistical differences ($p<0.05$) are marked by white dotted lines. Mice numbers: Male *Syngap1*$^{+/+}$ n=6; Male *Syngap1*$^{+/-}$ n=4; Female *Syngap1*$^{+/+}$ n=5; Female *Syngap1*$^{+/-}$ n=4.

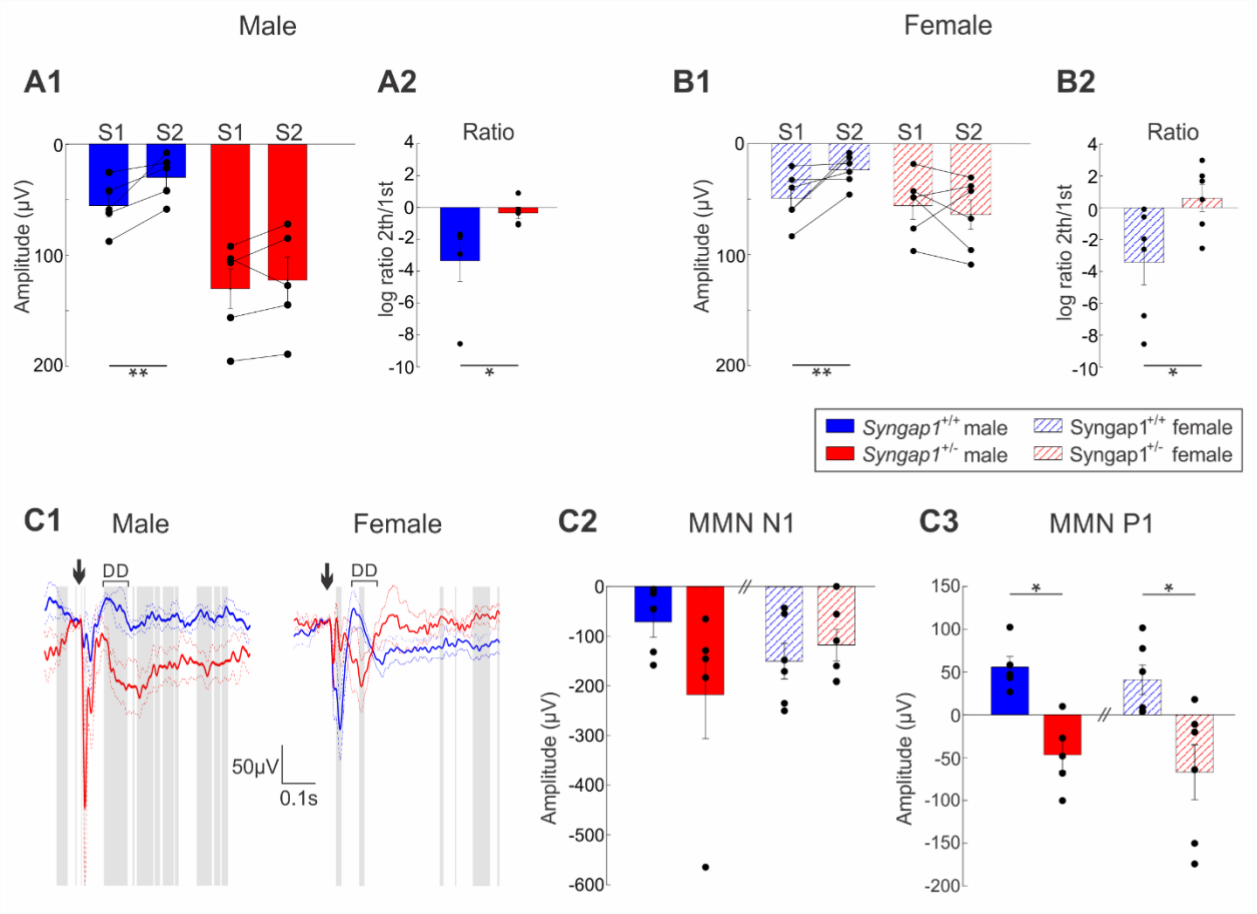


**Supplementary Figure 7. No sex differences are observed in habituation response and deviant detection in mice. A, B:** Responses to repetitive auditory stimuli in male and female mice. **A1, B2:** Bar plots show P1 amplitude after first (S1) vs second (S2) sounds in **(A1)** male (Syngap1$^{+/+}$, Wilcoxon signed rank, z=-2.0226, p=0.0431 and Syngap1$^{+/-}$, Wilcoxon signed rank, z=-0.6742, p=0.5002) and **(B1)** female (Syngap1$^{+/+}$, Wilcoxon signed rank, z=-2.2014, p=0.0277 and Syngap1$^{+/-}$, Wilcoxon signed rank, z=0.7338, p=0.4631) mice. **A2, B2**: Bar plots show the logarithmic ratio S2/S1 (two-way ANOVA; $F_{genotype}$ (1,18)=10.2093, p=0.0050, $F_{sex}$(1,18)=0.1472, p=0.7057, $F_{genotype*sex}$(1,18)=0.2098, p=0.6524) Tukey *posthoc* test shows genotyping effect in both males (A2, p=0.0455) and females (B2, p=0.0165). **C1-C3** Deviant detection in male and female mice. **C1**: superposed MMN (deviant - standard) traces from *Syngap1*$^{+/+}$ (blue) and *Syngap1*$^{+/-}$ (red) male and female mice. **C2:** Bar plots of MMN N1 component. Two-way ANOVA, $F_{genotype}$ (1,18)=1.2765, p=0.2734, $F_{sex}$(1,18)=0.0046, p=0.9464, $F_{genotype*sex}$(1,18)=3.1625, p=0.0922. Tukey *posthoc* test shows no genotype effect in males (p=0.2360) and females (p=0.9623). **C3,** Bar plots of MMN P1 component. Two-way ANOVA; $F_{genotype}$ (1,18)=21.7271, p=0.0002, $F_{sex}$(1,18)=0.5694, p=0.4603, $F_{genotype*sex}$(1,18)=0.0203, p=0.8882). Tukey *posthoc* test showed genotype effect in both males (p=0.0109) and females (p=0.0313). Shadow gray bars represent significant differences (p<0.05) between groups. Arrows indicate the start of stimulation. Mice numbers: Male *Syngap1*$^{+/+}$ n =5; Male *Syngap1*$^{+/-}$ : n=5; Female *Syngap1*$^{+/+}$ n=6; Female Syngap1+/- n=6.
